# MAPK dependent IL-33 responses define a conserved inflammatory programme in mast cells

**DOI:** 10.64898/2026.05.29.728623

**Authors:** Megan C. Sumoreeah, Iain R. Phair, Nicola J. Darling, J. Simon C. Arthur

## Abstract

Interleukin-33 (IL-33) is a key cytokine in mast cell mediated immunity, promoting inflammatory cytokine production without inducing degranulation. Here, we compared IL-33 induced proteomic responses across three mast cell culture systems, Foetal Liver derived Mast Cells (FLMCs), Bone Marrow derived Mast Cells (BMMCs), and Peritoneal Mast Cells (PMCs), using quantitative data-independent acquisition mass spectrometry.

Although baseline proteomes were largely conserved across all mast cell types, clear differences were observed between culture systems. PMCs exhibited a more mature phenotype, characterised by higher abundance of granule-associated proteins and lower levels of proteins involved in metabolism and translation. In contrast, FLMCs and BMMCs displayed higher levels of biosynthetic and metabolic machinery, consistent with a less differentiated state.

IL-33 stimulation induced a conserved proteomic programme across all mast cell types, enriched for inflammatory signalling pathways, cytokine production, and enzymes involved in prostaglandin and biogenic amine biosynthesis. Pathway analysis demonstrated robust activation of nuclear factor κB (NFκB) associated signalling, with a relative enrichment of components linked to non-canonical NFκB signalling and tumour necrosis factor (TNF) receptor associated pathways.

Mechanistically, IL-33 driven proteomic remodelling was strongly regulated by mitogen-activated protein kinase (MAPK) signalling. p38 MAPK emerged as the dominant regulator of the IL-33 response, with ERK1/2 contributing to a subset of induced proteins. These pathways differentially regulated key effector outputs, including IL-6, IL-9, IL-1 family cytokines, and enzymes required for prostaglandin, serotonin, and histamine biosynthesis.

Together, these data define conserved IL-33 dependent inflammatory programmes across mast cell differentiation states and demonstrate how MAPK signalling pathways shape the composition of mast cell effector responses.

## Introduction

Mast cells are tissue-resident granulocytes that are involved in host defence, allergic inflammation, and tissue homeostasis(1-3). They are characterised by dense cytoplasmic granules that contain a range of inflammatory mediators including mast cell proteases, proteoglycans and biogenic amines such as histamine and serotonin(4). Mast cells are predominately located at barrier sites such as the skin, lung, and gastrointestinal tract, where they can rapidly respond to infection, tissue damage and allergic stimuli(1-3). While classically associated with IgE-mediated allergic responses, mast cells are increasingly recognized as having multiple roles within innate and adaptive immunity(5-8), and this is facilitated by their expression of a range of immune receptors including Toll-like receptors (TLRs) and the Interleukin-33 (IL-33) receptor(9-12). Mast cell activation can take two main forms, degranulation, where there is a rapid release of preformed inflammatory agents from cytoplasmic granules, and *de novo* production of cytokines, prostaglandins and leukotrienes(4, 13, 14). Some stimuli, such as those that activate the IgE receptor initiate both of these processes, while others, such as TLR ligands or IL-33, do not on their own promote degranulation but do induce *de novo* cytokine production and release(10-12, 14-17).

IL-33 is a member of the IL-1 family of cytokines that functions as an alarmin following cellular stress or tissue damage. It is constitutively expressed and stored as a DNA-binding protein in epithelial and endothelial cells, and is released following necrosis or mechanical injury, thereby alerting the immune system to barrier disruption(18, 19). IL-33 signals through a heterodimeric receptor comprised of ST2 (encoded by Il1rl1) and the IL-1 receptor accessory protein (Il1rap), which activate downstream Myd88-dependent pathways that culminate in NFκB and MAPK signalling(20). *In vivo*, increased expression and release of IL-33 is detected in response to allergens, helminth infection and viral pathogens where it exerts broad immunomodulatory effects(21-23).

Mast cells express high levels of ST2, making them potentially one of the most responsive immune cell types to IL-33(24). IL-33 directly activates mast cells independently of IgE, promoting the production of cytokines including IL-13, IL-6 and TNF(10-12, 14-17). In contrast to other classical mast cell activating stimuli, IL-33 does not directly cause mast cell degranulation(11). It has however been reported to potentiate IgE mediated mast cell activation, lowering activation thresholds and amplifying degranulation and mediator release(25). In addition, co-stimulation of human skin mast cells with IL-33 and MAS-related G protein coupled receptor X2 (MRGPRX2) agonists amplifies both cytokine secretion and degranulation (26). These effects position IL-33 as a key regulator of mast cell effector function, linking tissue damage to rapid inflammatory responses.

Beyond acute activation, IL-33 has been implicated in shaping mast cell development, survival, and phenotypic plasticity. Emerging evidence suggests that IL-33 contributes to mast cell maturation within tissues and may influence their long-term functional programming, including changing their responsiveness to stimuli inducing degranulation and increasing their sensitivity to thymic stromal lymphopoietin (TSLP)(10, 27-29). IL-33 also promotes mast cell survival via upregulation of the pro-survival protein BCL-xL (Bcl2l1) and therefore may contribute to the persistence of mast cells within tissues during inflammation(30). However, the extent to which IL-33 drives distinct mast cell subsets or context-dependent responses remains incompletely understood. IL-33 and mast cell function have been implicated in a range of pathological conditions, including allergic asthma, atopic dermatitis, chronic urticaria, fibrosis and cardiovascular inflammation, underscoring their broad clinical relevance (5, 7, 8, 31, 32).

Mast cells are hard to study at a biochemical level. Their low numbers and location deep within tissue make the purification and culture of mast cells challenging. Mast cells are also a heterogenous cell type, as their final maturation occurs within tissues and is heavily influenced by the tissue environment(33). To circumvent this, several methods have been described to differentiate mast cells in culture from either the foetal liver or bone marrow. While these yield high numbers of cells, they do not have all the characteristics of mature, tissue-resident mast cells, particularly with respect to the formation of mature mast cell granules(34). Mast cells can also be cultured from the peritoneal cavity and while these may represent a more mature form of mast cells, the numbers obtained are much lower than for other mast cell culture methods(35). Despite their limitations, mast cell cultures have been informative in dissecting the molecular details of stimulus-specific intracellular signalling responses.

In this study we use data independent acquisition (DIA) based proteomic methods to compare 3 commonly used mast cell culture methods, Foetal Liver-derived Mast Cells (FLMCs), Bone Marrow-derived Mast Cells (BMMCs) and Peritoneal Mast Cells (PMCs) and characterise their response to IL-33. Using these methods we have compared different mast cell culture systems and identified a core set of proteins induced by IL-33. Using small molecule inhibitors, we also show that the activation of p38 MAPK by IL-33 is important for the induction of a subset of these proteins.

## Results

### Comparison of mast cell proteomes and their response to IL-33

To compare the responses of different mast cell culture systems to IL-33 stimulation, we used 3 different protocols; Foetal Liver-derived Mast Cells (FLMC), Bone Marrow-derived Mast Cells (BMMC) and Peritoneal Mast Cells (PMC). The proteomes of these cells were then determined with or without IL-33 stimulation for 24h using DIA mass spectrometry. 5 replicates were used for FLMCs and 4 for BMMCs and PMCs. Estimation of the total cellular protein mass showed that unstimulated FLMC and BMMCs have similar protein levels but that PMCs had a slightly higher protein content (Figure 1A). In all 3 cell types, IL-33 stimulation resulted in a small increase in cellular protein levels. In total 8624 protein species were identified across the different conditions, however of these 365 were discarded as they were detected in less than 3 replicates across the mast cell types analysed. Mast cells are frequently identified by the expression of the stem cell factor (SCF) receptor (Kit or CD117) and the high affinity IgE receptor (FcepsilonR1)(1, 2). The proteomics identifies similar levels of both Kit and the alpha subunit of FcepsilonR1 (Fcer1a) in FLMCs, BMMCs and PMCs (Supplementary figure 1A, B). The other two subunits of the IgE receptor (Fcer1b/Ms4a2 and Fcer1g) were also found in all 3 cell types, although levels of these were lower in BMMCs and PMCs relative to FLMCs (Supplementary figure 1C, D). Mast cells are also responsive to IL-33(10-12, 14-16), and in line with this, all 3 cell types also expressed the IL-33 receptor subunit (Il1rl1, ST2) as well as the IL-1 receptor accessory protein (Il1rap), although the Il1rl1 chain was present at much higher levels (Supplementary figure 1E, F). Recently mast cells have been found to degranulate in response to stimulation with some secretagogues and neuropeptides which activate Mrgprb2 (MrgprX2 in humans)(36, 37). Mrgprb2 was not detected in BMMCs in this dataset, but was found in FLMCs and PMCs, with the highest levels in PMCs (Supplementary figure 1G).

**Figure 1.**
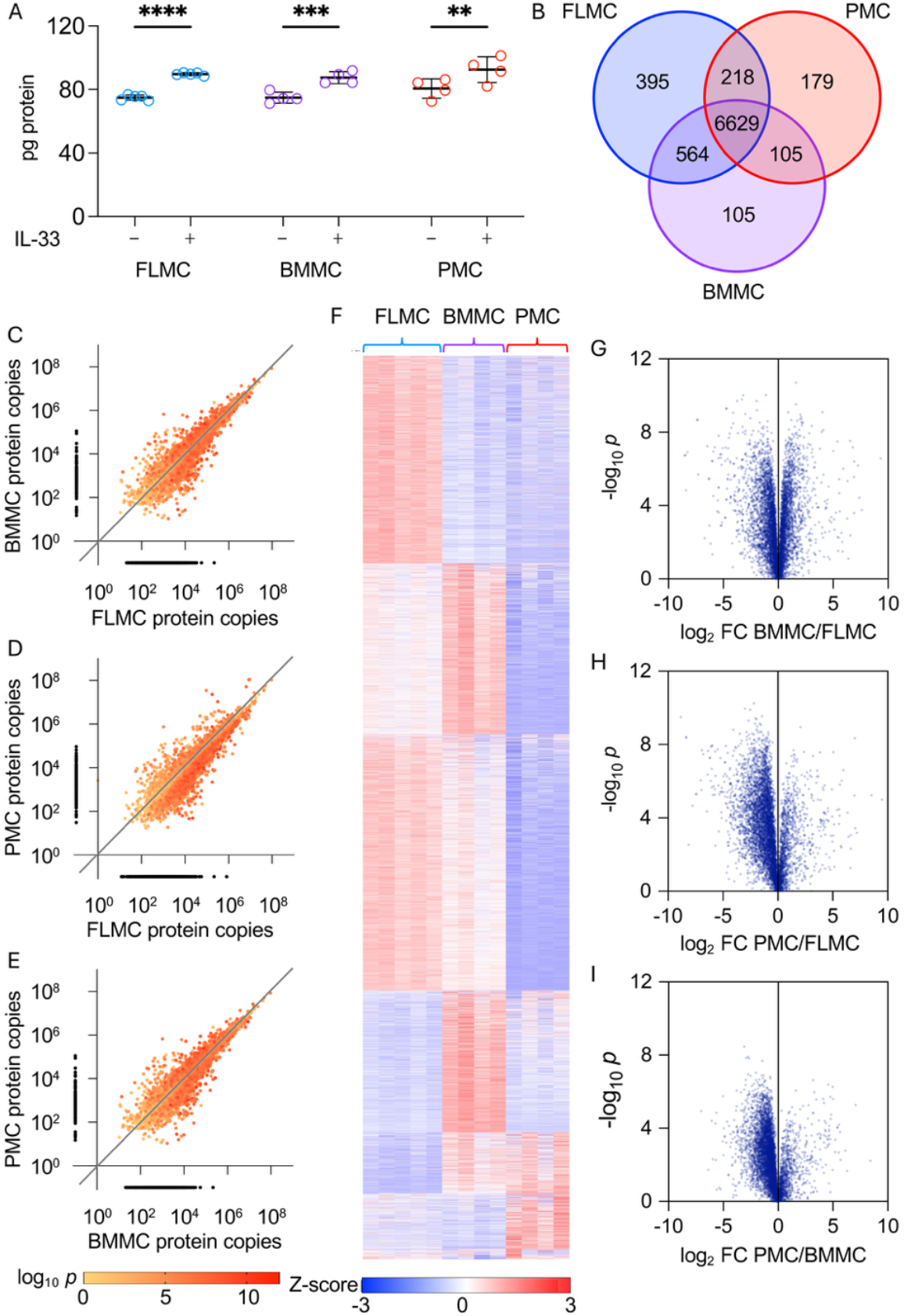
Proteomic characterisation of FLMC, BMMC and PMCs. DIA-based mass spectrometry was used to characterise the proteomes of FLMCs (n=5), BMMCs (n=4) and PMCs (n=4) with or without IL-33 stimulation for 24 h. (A) Estimated protein content per cell. (B) Numbers and overlap of proteins identified in each mast cell type. (C–E) Pairwise comparison of estimated protein copy numbers between the indicated unstimulated mast cell types. The levels of proteins found in only one of the cell types are indicated by the black symbols outside of the axes. (F) Heat map showing protein levels between the unstimulated cell types with k-means clustering into 6 groups. (G–I) Volcano plots showing pairwise comparisons between the indicated unstimulated mast cell types. In A, for comparisons between the control and IL-33 conditions, *p*<0.01 is indicated by **, <0.001 by *** and <0.0001 by **** (Two-way ANOVA, cell type F=3.277 p=0.0587, treatment F=54.662 p<0.0001, interaction F=0.2656p=0.7694 and Tukey’s post hoc tests).

In the unstimulated mast cells, 7806 proteins were detected in the FLMCs, 7403 for the BMMCs and 7131 for the PMCs with 6629 of these proteins being found in all 3 mast cell types (Figure 1B). For proteins expressed in more than one cell type there was a good correlation in the levels between two different cell types (Spearman coefficient of 0.9242 for FLMC vs BMMC, 0.8991 for FLMC vs PMC and 0.9329 for BMMC vs PMC, Figure 1C-E). Despite these correlations, there were clear groups of proteins showing higher expression in specific mast cell culture types (Figure 1F), as well as a clear trend for many proteins being present at slightly lower levels in PMC than either FLMCs or BMMCs (Figure 1F-I). This could be accounted for by FLMCs and BMMCs representing a less mature form of mast cells than PMCs, with fewer mature granules.

In mice, mast cells have been classed into mucosal and connective tissue mast cells, based on a combination of location and granule content. The use of scRNAseq has recently expanded on this concept, identifying a list of 195 differentially regulated genes between MrgprB2^pos^ connective tissue mast cells and MrgprB2^neg^ mucosal mast cells (33). 159 of these were identified in the 3 proteomic data sets reported here, with FLMC and BMMCs expressing higher levels of proteins associated with mucosal mast cells. Analysis of correlations between replicates for this protein subset indicated that FLMCs and BMMCs were closely associated with each other, with PMCs forming a divergent group (Figure 2A, Supplementary figure 2).

**Figure 2.**
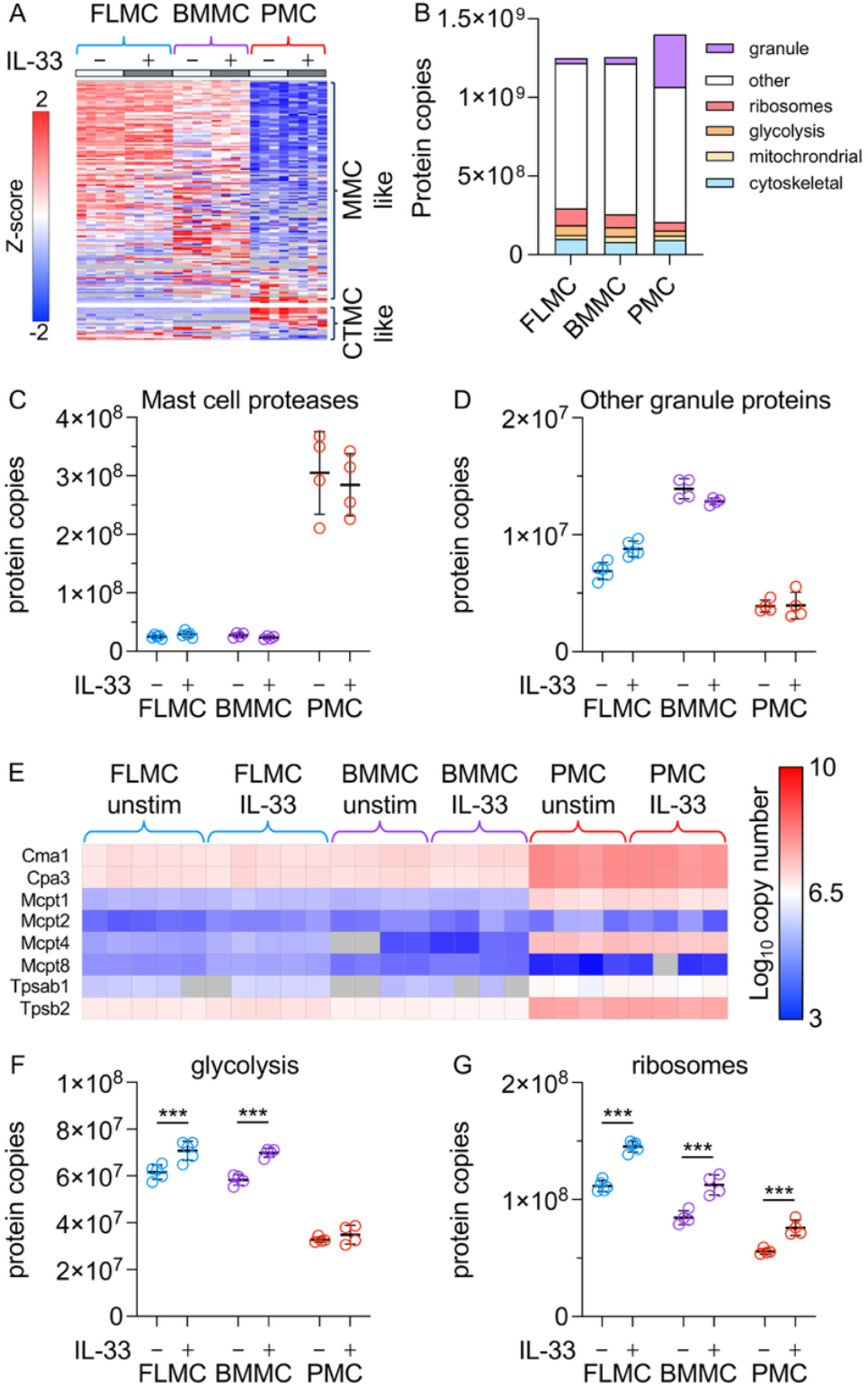
Analysis of protein subgroups in cultured mast cells. (A) A set of 159 proteins previously found to be differentially regulated between mucosal and connective tissue mast cells was extracted from the FLMC, BMMC and PMC proteomic data. Heat map shows z-score of protein concentrations across the 26 replicates. (B-D) Using data from the proteomics experiment shown in Figure 1, copy numbers for proteins involved in the cytoskeleton, mitochondrial function, glycolysis, ribosome, mast cell-specific proteases and other granule-related proteins were calculated. (B) Copy numbers of the indicated protein groups as a proportion of total protein copies in unstimulated FLMCs, BMMCs and PMCs. (C) Total copy number for mast cell-specific proteases in FLMCs, BMMCs and PMCs with or without IL-33 stimulation for 24 h. (D) As (C) but combined copy number of other granule proteins. (E) Heat map showing the copy numbers of the individual mast cell-specific proteases detected in the proteomics. (F) As (C) but combined copy number for proteins involved in glycolysis. (G) As (C) but combined copy number of ribosomal proteins and proteins involved in ribosome biogenesis. For C, D, F and G, lines show mean with standard deviation. For comparisons between unstimulated and IL-33 stimulated conditions, *p*<0.05 is indicated by *, <0.01 by ** and <0.001 by *** (Two-way ANOVA with Sidak’s post hoc tests). ANOVA F and p values are given in Supplementary Table 5. In heatmaps, grey indicates replicate where a protein was not detected.

To examine granule content, the levels of proteins known to be contained in mast cell granules (based on(4)) was determined. PMCs expressed much higher levels of granule proteins compared to FLMC or BMMCs, with approximately 24% of proteins in PMCs accounted for by granule proteins compared to 3% for FLMCs and BMMCs (Figure 2B). This was predominantly due to increased levels of mast cell specific proteases in PMCs relative to the other two mast cell culture types (Figure 2C-E). An exception to this was Mcpt8 (Figure 2E), which was lower in PMCs. Of note, this protein has been previously associated with basophils rather than mast cells (38), and its expression in FLMCs and BMMCs may indicate these cells are not fully differentiated mast cells. Mast cell specific proteases fall into 3 groups, tryptases, chymases and carboxypeptidases. The ratios of these three protease types were similar between BMMCs and PMCs, although FLMCs had a slightly lower chymase : tryptase ratio (Supplementary Figure 3A).

In contrast to mast cell proteases, differences in other granule proteins across FLMCs, BMMCs and PMCs were less dramatic. β-hexosaminidase is found in mast cell granules and has frequently been used as a marker for degranulation, and may target bacterial cell walls following its release(39). BMMCs and PMCs expressed similar levels of β-hexosaminidase with lower levels found in FLMCs (Supplementary figure 3B, C). Mast cell granules also express arylsulfatases(40). Levels of arylsulfatases were also higher in BMMCs and PMCs than FLMCs (Supplementary Figure 3D, E). Cathepsins were also identified in all 3 mast cell types, with slightly higher levels in BMMCs (Supplementary Figure 3F). Granzymes are a group of proteases more commonly associated with NK and T cell secretory granules(41), although they have been reported to be expressed in mast cells(42, 43). FLMCs and BMMCs were also found to express high levels of granzyme B (Gzmb), whereas very little granzyme B was found in PMCs (Supplementary Figure 3G). Overall IL-33 did not promote an increase in granule proteins in any of the 3 cell types, suggesting that a major function of IL-33 is not to promote granule formation. One exception to this was serglycin, a protein implicated in the storage of mast cell specific proteases(44, 45). Serglycin levels were increased by IL-33 treatment in FLMCs and BMMCs (Supplementary figure 3H). Unexpectedly despite their higher levels of mast cell proteases, PMCs had the lowest levels of serglycin of the 3 cell types and this was not affected by IL-33.

While they expressed lower levels of mast cell proteases, the proportion of the proteome comprising enzymes involved in glycolysis was higher in FLMCs and BMMCs than PMCs (Figure 2F). FLMCs and BMMCs also had higher levels of ribosomal proteins and proteins involved in ribosome biogenesis (Figure 2G). In all 3 mast cell types, IL-33 stimulation significantly increased the levels of these proteins, which would be consistent with IL-33 increasing the protein content of the cells (Figure 1A).

### Identification of a core set of IL-33 induced proteins in mast cells

IL-33 is well established to stimulate the *de novo* production of several cytokines, including TNF, IL-6, IL-9 and IL-13 without causing degranulation(10-12, 14-16, 46). While the majority of the cytokine produced may be secreted, IL-6, IL-9 and IL-13 were consistently detected in the cell lysates from IL-33 stimulated cells. In contrast they were absent in the unstimulated condition for FLMCs, BMMCs and PMCs (Figure 3A-C). TNF was detected in lysates from IL-33 stimulated FLMCs, but not BMMCs or PMCs (Figure 3D). All 3 types of mast cell were also found to induce both IL-1α and IL-1β after IL-33 stimulation (Figure 3E, F).

**Figure 3.**
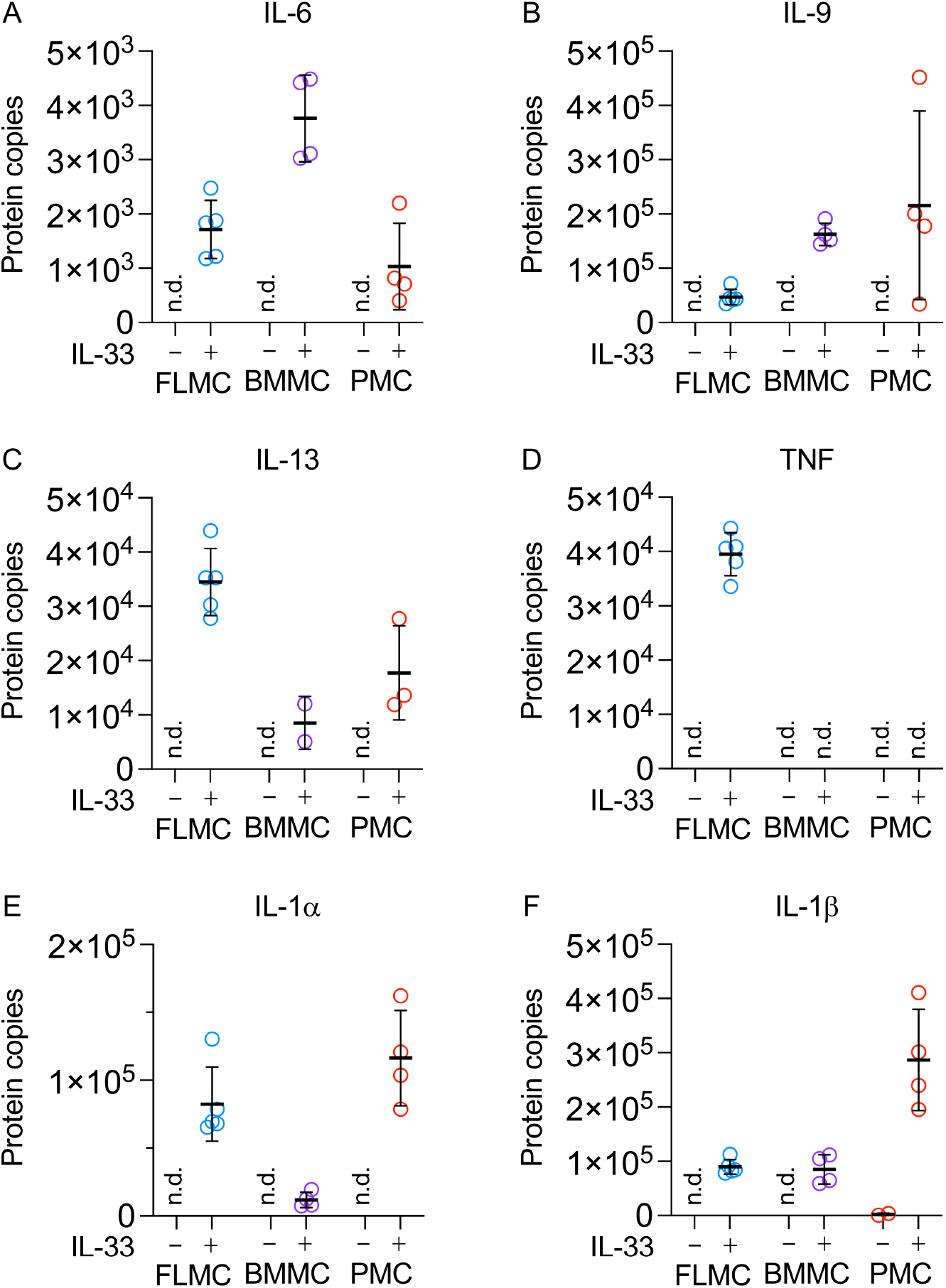
Cytokine levels in FLMCs, BMMCs and PMCs. Copy numbers of IL-6 (A), IL-9 (B), IL-13 (C), TNF (D), IL-1α (E) and IL-1β (F) estimated from the proteomic data in FLMCs, BMMCs and PMCs with or without 10 ng/mL IL-33 stimulation for 24 h. Plots show mean and standard deviation with individual biological replicates as symbols. n.d. indicates not detected.

To look at effects of IL-33 on protein abundance, we defined groups of up and down regulated proteins as those with log2 fold change of more than 2 standard deviations away from the median and a *p* value of less than 0.01. In addition, a protein was regarded as regulated if it was present in all of the replicates for one condition but none of the other. For this analysis estimated protein concentrations, rather than copy number, was used to allow for a focus on proteins whose change did not scale with the increase in cell size following IL-33 stimulation. Using this approach, 99 proteins were upregulated in FLMCs along with 35 more that were present in all the IL-33 stimulated replicates but none of the unstimulated ones. 87 proteins were downregulated, with a further 6 present in all of the unstimulated replicates but none of the stimulated ones (Figure 4A, supplementary table 1). 137 proteins were upregulated in BMMCs with a further 62 present in all stimulated but no unstimulated cells while 44 were downregulated and a further 39 present in all unstimulated but no unstimulated cells (Figure 4B, supplementary table 1). For PMCs, 90 proteins were upregulated, with a further 31 present in all stimulated but no unstimulated cells, while 16 were downregulated and a further 22 present in all unstimulated by no unstimulated cells (Figure 4C, supplementary table 1). For proteins regulated in FLMCs there was a reasonable correlation between their fold change in BMMCs and PMCs (Spearman coefficients of 0.774 and 0.608 respectively, however a subset of proteins that were regulated in FLMCs did not show significant regulation in BMMCs and PMCs (Figure 4D, E). Similar results were found for proteins regulated in BMMCs and PMCs (Figure 4F-I).

**Figure 4.**
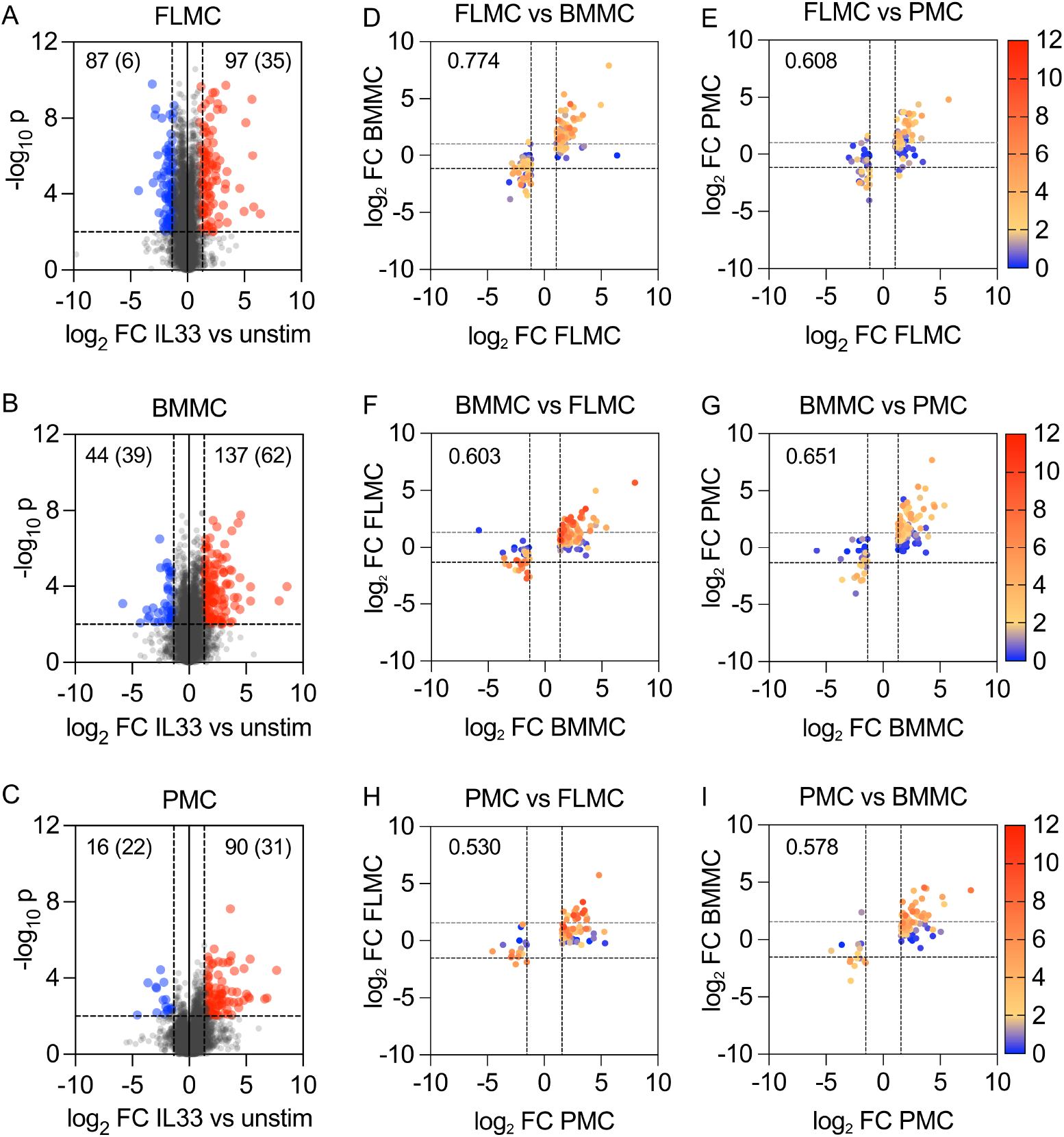
IL-33-stimulated proteome remodelling in mast cell cultures. FLMCs, BMMCs or PMCs were left unstimulated or treated with 10 ng/mL IL-33 for 24 h, and protein expression determined by DIA-based mass spectrometry, with protein concentrations estimated using the histone ruler method. Regulated proteins were defined as having a log2 fold change (FC) greater than 2 standard deviations away from the median and *p* < 0.01. (A–C) Volcano plots showing the effect of IL-33 in FLMCs (A), BMMCs (B) and PMCs (C). Red indicates upregulated proteins and blue downregulated. Numbers indicate the number of proteins identified, with the number in brackets showing the number of proteins found in all biological replicates for one condition but absent in all replicates for the other condition. (D–I) Plots showing the fold change for proteins identified as regulated in one cell type (x-axis) plotted against fold change in a second cell type (y-axis), with colour indicating the -log10 *p-*values for the change in the second cell type.

Of the upregulated proteins that met the cutoffs used, 38 fell into this category in all 3 cell types analysed (Figure 5A), including IL-6 and IL-9. IL-33 also increased the levels of a number of signalling proteins and transcription factors as well as enzymes important in mast cell functions such as cyclooxygenase-2 (Ptgs2), tryptophan hydroxylase 1 (Tph1) and histidine decarboxylase (Hdc) (Figure 5B). In contrast, 47, 93 and 54 proteins only met the cutoff in FLMCs, BMMCs and PMCs respectively (Figure 5A). However, comparing across cell types these proteins may have been affected by IL-33 in more than one cell type, but only met the fold change and significance cutoffs in 1 (Figure 5C, Supplementary table 1). While there was clear overlap in proteins upregulated by IL-33 in all 3 mast cell types, there was less apparent overlap in the downregulated proteins (Supplementary figure 4). Enrichment analysis (using FDR < 0.05) for downregulated proteins did not highlight any process for BMMCs and PMCs, although there was an enrichment for proteins involved in the cell cycle and cell division in FLMCs (Supplementary table 2). Enrichment analysis of upregulated proteins against the GO term Biological Process and KEGG databases returned terms relating to inflammation, the immune system and signalling pathways known to control innate immune cells (Supplementary Table 2). Eight KEGG pathways and 5 GO terms were found in all 3 mast cell types (Figure 5D).

**Figure 5.**
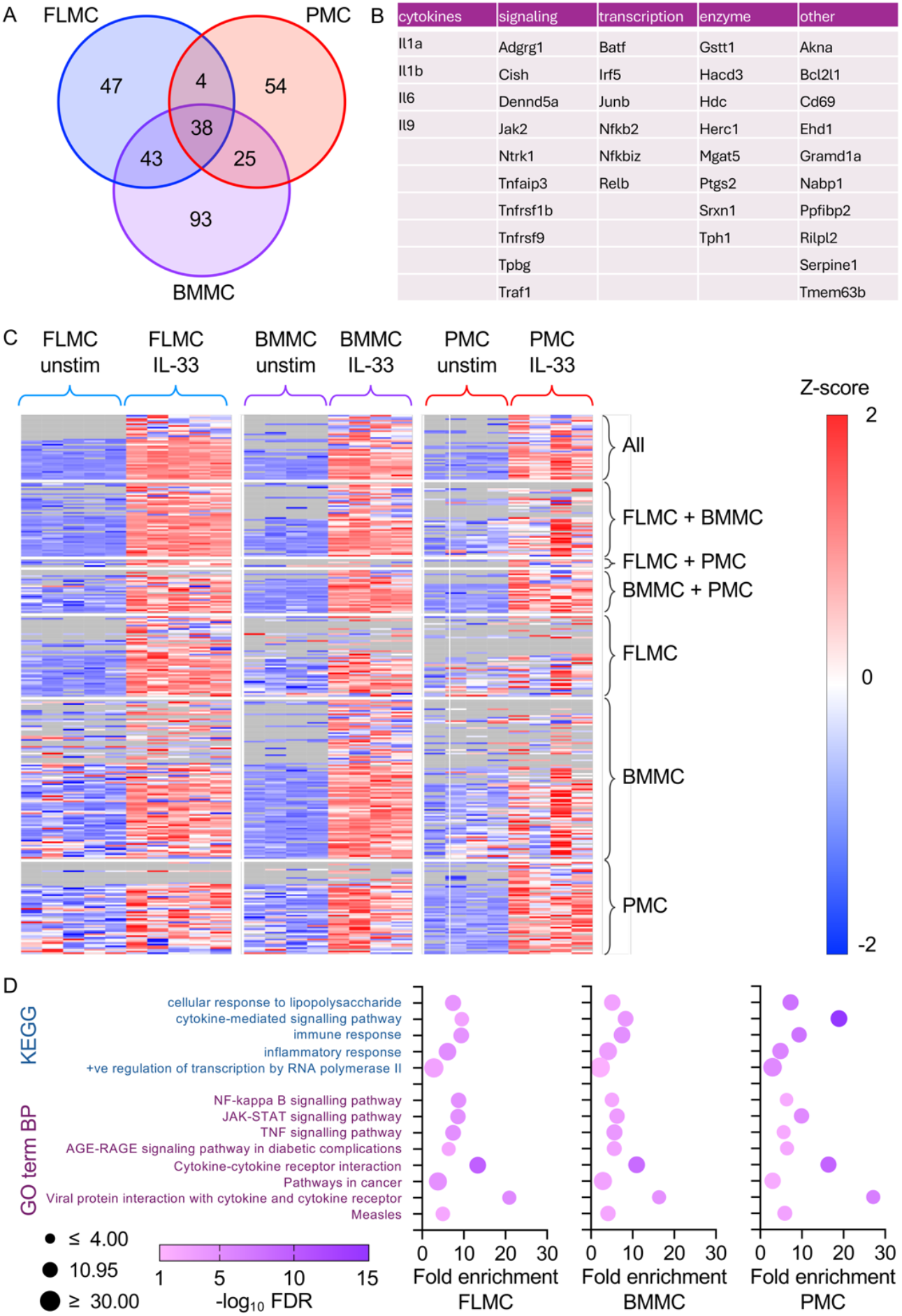
IL-33 upregulates a core set of proteins across different mast cell culture types. (A) Intersects between the proteins identified as upregulated in FLMCs, BMMCs and PMCs (based on a log2 fold change greater than 2 standard deviations from the median and *p* < 0.01 or present in all IL-33 replicates but no unstimulated replicates). (B) Proteins identified as upregulated in all 3 mast cell types categorised into cytokines, signalling proteins, transcriptional regulators, enzymes and other protein classes. (C) Heat map of IL-33-regulated proteins in FLMCs, BMMCs and PMCs. Grey indicates replicate where a protein was not identified. (D) Dot plots showing the enriched terms found for all 3 mast cell types for upregulated proteins against the KEGG and GO term Biological Process databases.

One pathway that was highlighted in all 3 mast cell types was the NFκB pathway, which is known to be activated downstream of Myd88-dependent signalling(47). In line with this, analysis of the gene promoters for IL-33 upregulated proteins also showed an enrichment for NFκB binding sites (Supplementary figure 5). Activation of NFκB has been broadly characterised into canonical and non-canonical pathways(48, 49). The NFκB family is made up of 5 proteins, c-Rel, RelA, RelB, p105 (encoded by Nfkb1 and which is processed to p50) and p100 (Nfkb2, which is processed to p52). Canonical NFκB typically involves the c-Rel, RelA and p105/p50 NFκB proteins, while non-canonical activation is more closely associated with RelB and p100/p52. While canonical activation is the main pathway activated acutely downstream of IL-33 signalling(16, 17, 26, 50), it was the subunits more closely associated with non-canonical activation that were more strongly upregulated by IL-33 (Figure 6 A-E). Activation of the non-canonical pathway requires the phosphorylation of p100 by IκB kinase (IKK)α, which promotes its processing to p52. IKKα is in turn activated by NIK, which in resting cells is normally ubiquitinated and degraded. Non-canonical NFκB activation occurs downstream of a subset of TNF receptor family members(48, 49). In the presence of activating ligands these receptors interact with Traf2, Traf3 and potentially Traf1 and Traf5, which then allows for the stabilisation of the NIK protein and activation of IKKα as well as activation of the canonical NFκB pathway. Traf1, Traf2 and Traf3 were also found to be upregulated following IL-33 stimulation (Figure 6F-I), although NIK was not detected. Mining the proteomic data for the presence of TNF family receptors and TNF family ligands revealed the presence members of these families (Figure 6J). TNF family receptors can be divided into those that contain a death domain and those which do not(51). Receptors lacking the death domain interact directly with Traf proteins to activate the canonical and/or non-canonical NFκB pathways. Two TNF receptor family members lacking the death domain with the potential to activate the non-canonical pathway, Tnfrsf1b (TNFR2) and Tnfrsf9 (CD137) were detected in all 3 mast cell types tested, and were also upregulated by IL-33 (Figure 6J). In addition, FLMC also upregulated a third non death domain containing receptor, Tnfrsf8 (CD30, Figure 6J). Mast cells are known to express TNF, the ligand for Tnfrsf1b, following IL-33 stimulation and the proteomic data also found that FLMCs constitutively expressed Tnfsf8 and Tnfsf9, the ligands for Tnfrsf8 and Tnfrsf9 respectively (Figure 6J). IL-33 also upregulated Gramd1a, a protein involved in cholesterol sensing and related to there was also a trend for IL-33 to increase key enzymes in cholesterol metabolism including Hmgcr, Sqle and Ldlr (Supplementary figure 6).

**Figure 6.**
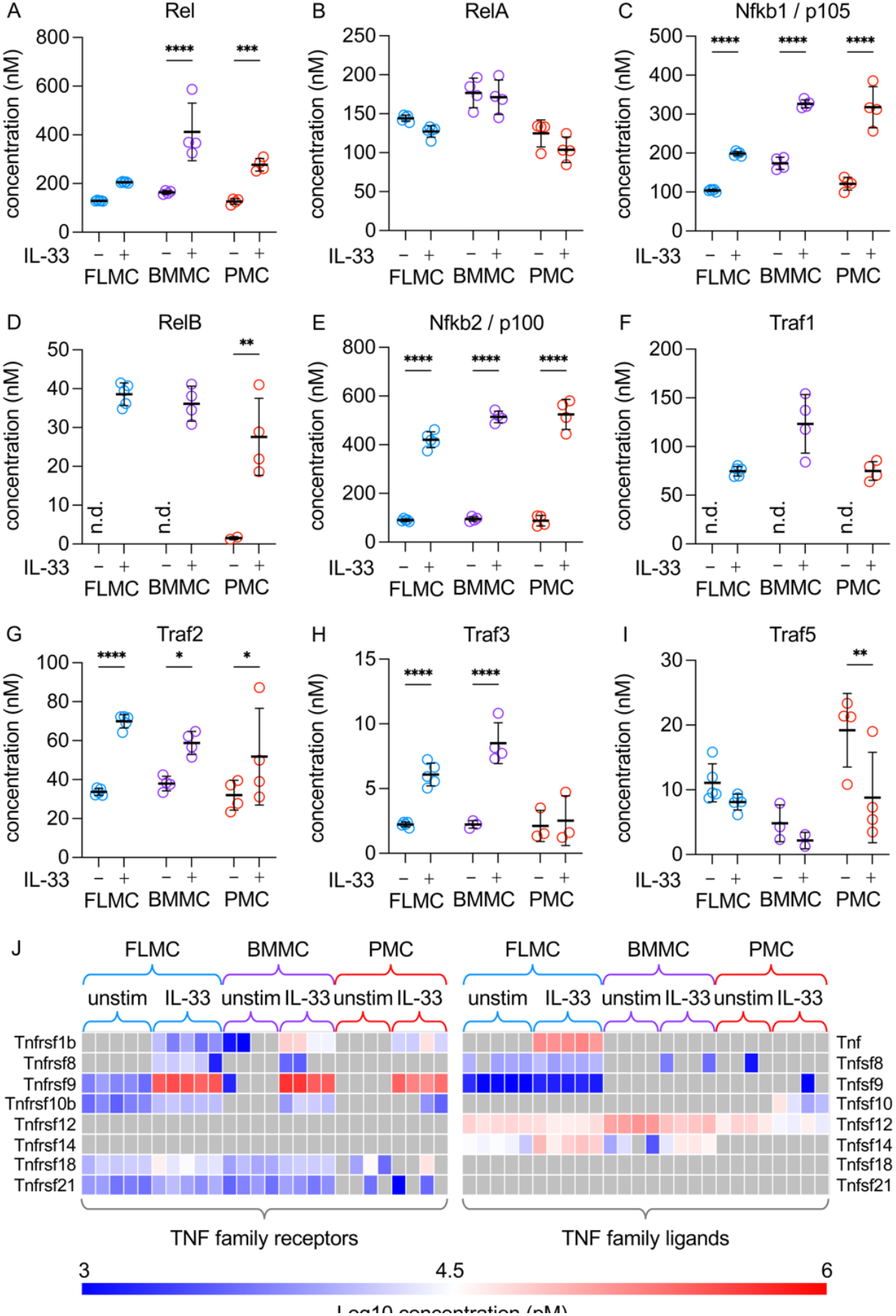
Abundance of NFκB components in mast cells. FLMCs, BMMCs or PMCs were left unstimulated or treated with 10 ng/mL IL-33 for 24 h, and protein abundance was analysed by proteomics. (A–I) The estimated concentrations of the indicated proteins in the different mast cell types are shown. Graphs show mean and standard deviation with individual biological replicates shown by symbols. n.d. indicates a condition where the proteins were not identified. For comparisons between unstimulated and IL-33 stimulated conditions, p<0.05 is indicated by *, p<0.01 by **, p<0.001 by *** and p<0.0001 by **** (Two-way ANOVA and Sidak’s multiple comparison tests). F and p values for the tests are given in Supplementary Table 5. (J) Heat map showing the log10-transformed concentrations for members of the TNF receptor and TNF ligand families. Grey squares indicate a protein was not detected.

### p38 MAPK regulates a subset of IL-33 induced proteins

Both the p38 and ERK1/2 MAPK pathways have previously been shown to be important in the regulation of cytokine production in mast cells (15, 16). To investigate how they may affect the proteomic response, BMMCs were treated with VX745, a p38α/β inhibitor, or PD184352, a MEK1/2 inhibitor that blocks the activation of ERK1/2(52, 53). Focusing on the core set of 38 proteins upregulated by IL-33 in all 3 mast cell types used, PD184352 treatment resulted in a significant decrease (*p*<0.01) in the induction of 10 proteins. VX745 had a greater effect, significantly reducing 20 proteins, while combination of VX745 and PD184352 had a greater effect than VX745 alone (Figure 7A). Looking more widely at all IL-33 induced proteins in BMMCs a similar trend was observed, with VX745 having a greater effect than PD184352 and the combination of VX745 and PD184252 having the greatest effect (Figure 7B-D, supplementary table 3). Strikingly, most IL-33 induced proteins showed a trend for a reduction in the presence of VX745, although in many cases this was small (less than two-fold) and only reached significance (*p*<0.01) for 43% of the IL-33 upregulated proteins.

**Figure 7.**
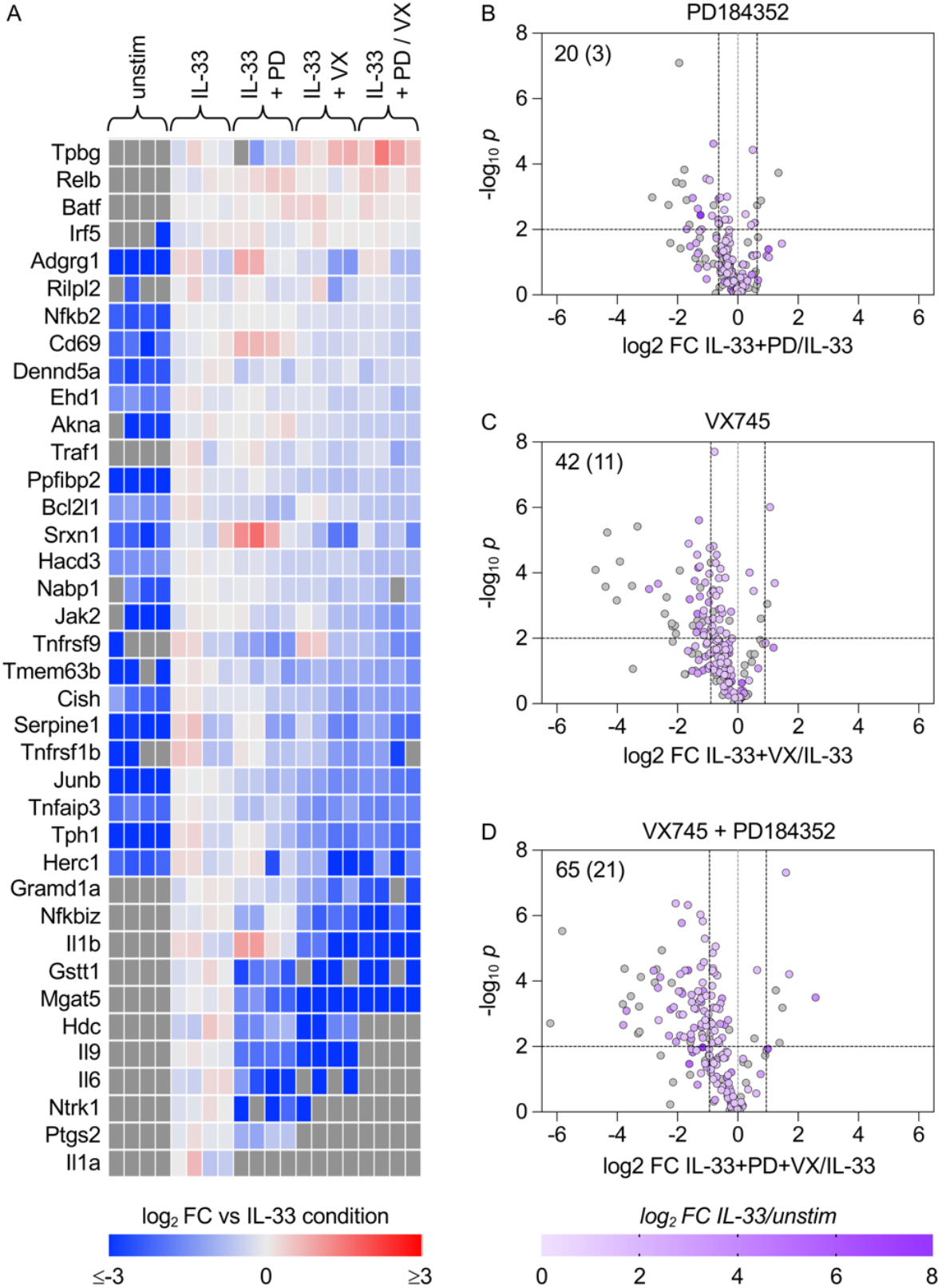
Role of MAPKs in regulating protein expression in response to IL-33. BMMCs were treated with either 2 μM PD184352 (PD), 1 μM VX745 (VX) or a combination of both inhibitors for 1h, as indicated, and the cells were then stimulated for 24 h with 10 ng/mL IL-33. Proteomes were then determined by DIA-based mass spectrometry. (A) Heat map showing the effect of PD184352 and VX745 on the subset of proteins identified as regulated by IL-33 in FLMCs, BMMCs and PMCs, as shown in Figure 6. (B–D) Volcano plots showing the effect of PD184352 (B), VX745 (C) or a combination of PD184352 and VX745 (D) compared to IL-33 alone specifically on the subset of 199 proteins found to be upregulated by IL-33 in BMMCs. Numbers in the top left quadrant indicate the number of proteins for which a log2 fold change is more than 1 standard deviation away from 0 and *p* < 0.01, with the number in brackets indicating proteins which were detected in none of the inhibitor-treated replicates.

In line with previous reports, IL-6 induction in response to IL-33 was reduced by inhibition of either ERK1/2 or p38, with a similar effect seen for IL-9 and IL-1α. However, IL-1β induction was only sensitive to p38 inhibition (Figure 8A-D). Mast cells are a potential source of prostaglandins (12, 50), and expression of the rate limiting enzyme in the production of these lipids, Ptgs2, was increased by IL-33. This increase was reduced by inhibition of either ERK1/2 or p38 (Figure 8E). Mast cells also produce serotonin and histamine, whose production is catalysed by Tph1 and Hdc respectively(54, 55). Both of these were increased following IL-33 treatment. Tph1 was only slightly reduced by PD184352 while the p38 inhibitor VX745 had a greater effect (Figure 8F). In contrast, induction of Hdc was sensitive to both ERK1/2 and p38 inhibition (Figure 8G). p38 is known to regulate protein abundance via various mechanisms including control of transcription, mRNA stability and translation(56-58). In mast cells roles for the mRNA binding proteins Zfp36l1 (also known as BRF1 or TIS11b) and Zfp36l2 (also known as BRF2 or TIS11d) have been proposed(15). These proteins bind to AU rich elements in the 3’ untranslated region (UTR) of target mRNAs in order to regulate their translation or stability. Both Zfp36l1 and Zfp36l2 were detected in the proteomic experiments, with Zfp36l1 being upregulated in a MAPK-dependent manner. Consistent with this, both Zfp36l1 and Zfp36l2 were detected in the proteomics, with Zfp36l1 found in the IL-33 treated cells only (Supplementary figure 7). Comparison of IL-33 regulated genes with a database of AU-rich containing mRNAs (59) showed that IL-33 upregulated proteins were more likely to contain an AU-rich element in their mRNA than IL-33 downregulated proteins (Figure 8H). Despite this there was no clear correlation between the size of the effect of VX745 on a protein and the presence of a potential AU-rich element in its corresponding mRNA (Figure 8I).

**Figure 8.**
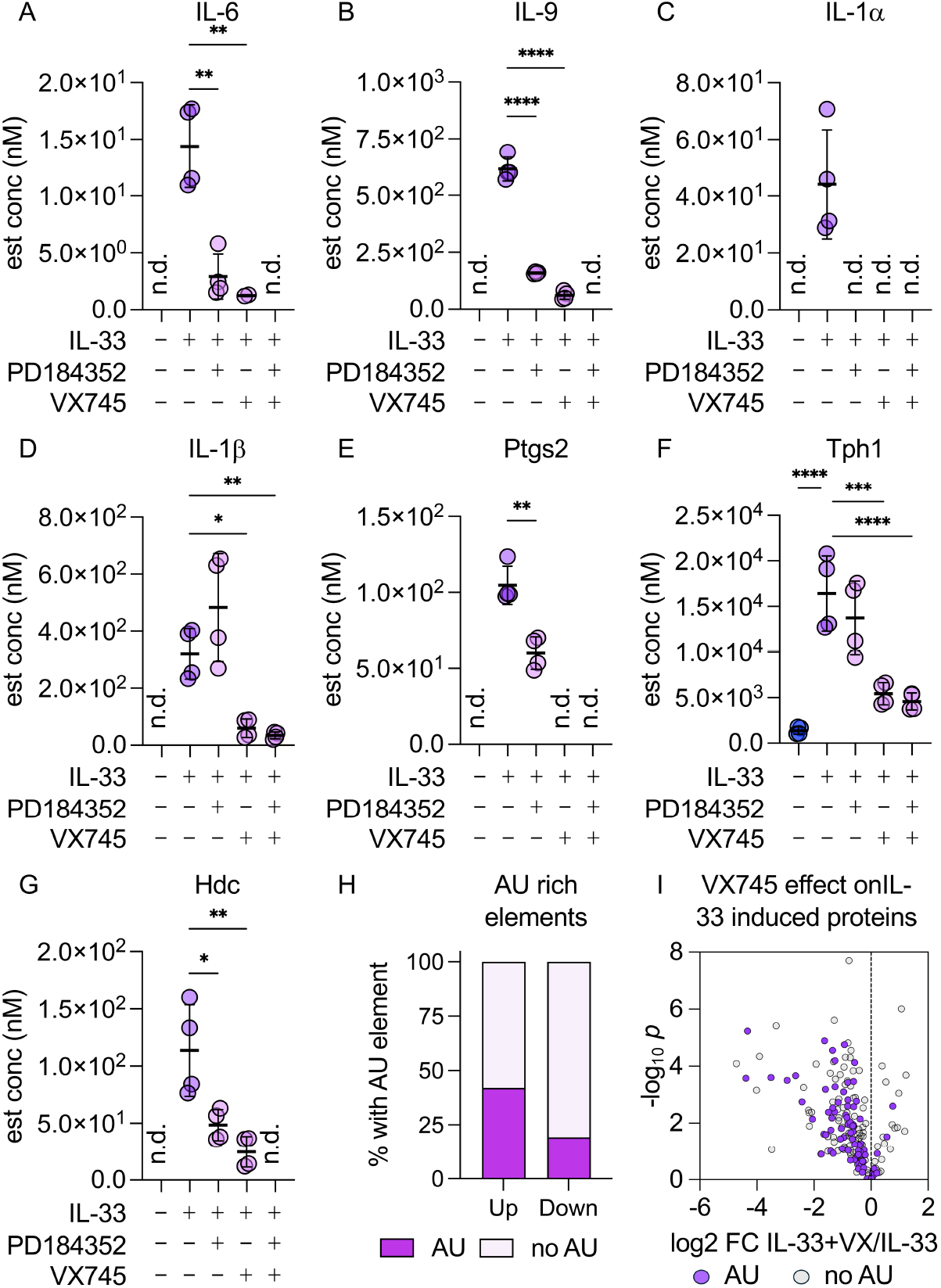
Effect of p38 inhibition of selected proteins in BMMCs. BMMCs were treated with 2 μM PD184352, 1 μM VX745 or a combination of both inhibitors before stimulation with IL-33 as described in Figure 6. (A-G) Plots show the estimated concentrations for IL-6 (A), IL-9 (B), IL-1a (C), IL-1b (D), Ptgs2 (E), Tph1 (F) and Hdc (G). Plots show mean and standard deviation with individual replicates as symbols. n.d. indicates not detected. For comparisons to the IL-33 alone condition *p*<0.05 is indicated by *, *p*<0.01 by **, *p*<0.001 by *** and *p*<0.0001 by **** (A, B, D, F, G -One ANOVA and Dunnett’s multiple comparison tests, E – two way Student’s t-test). F and p values for the ANOVA are given in supplementary table 5. (H) Percentage of proteins up or down regulated by IL-33 in BMMCs that contain potential AU rich elements in the 3’ UTR of their mRNAs based on information in the ARED AU Rich Element Database (https://brp.kfshrc.edu.sa/ared/#page-top). (I) Volcano plot showing the effect of VX745 (VX) on IL-33 upregulated proteins with those that contain a potential AU rich element in the 3’ UTR of their mRNA shown by purple symbols.p

## Discussion

Due to difficulties in isolating mast cells in sufficient numbers from tissue, various methods have been used to differentiate mast cells in culture in order to study them, especially with regards to the intracellular signalling processes that control mast cell function. We report here the proteomic characterisation of 3 mast cell culture systems from mice. All 3 systems expressed key mast cell markers, such as Kit and the high affinity IgE receptor. A proteomic characterisation of purified peritoneal mast cells has been reported previously, which identified 4292 proteins (60). The dataset here captured 88% of these proteins (based on gene names) but also expanded the identified number of proteins by around 3000. While an improvement, this is unlikely to represent the full proteome of these cells. Plum *et al* (60) reported that they could not detect innate immune receptors such as TLRs in their proteome, a result consistent with a second study (61). Despite this, mast cells are well established to respond to LPS stimulation (9), suggesting that these receptors while present are expressed at low levels. While we did identify TLR4 in some replicates, it was in the lowest 1% of proteins ranked by abundance suggesting that further proteomic depth will be required to fully elucidate which innate immune receptors are expressed in mast cells.

In vivo, bone marrow-derived hematopoietic stem cells mainly replenish the mucosal mast cell population (62), but following differentiation in culture with IL-3 these are often used to model connective tissue type mast cells (34). All 3 mast cell culture systems tested had similar ratios of chymases, tryptases and carboxypeptidases, with a low expression of Mcpt1 and Mcpt2. This profile would be consistent with a connective tissue rather than mucosal mast cell phenotype (63). More recently scRNAseq data has been used to define mast cell subtypes (33), comparing our proteomics results to this data, BMMC and FLMC showed higher abundance than PMCs for proteins associated with mucosal mast cells. Of note however was that mast cell proteases were found at much lower levels in BMMCs and FLMCs than PMCs. This would also be consistent with a less mature phenotype in BMMCs and FLMCs and in line with reports finding that these cells do not have high levels of mature granules (34, 64).

We identified a core set of 39 proteins that were strongly induced by IL-33 in all 3 mast cell culture systems tested. Similar to TLRs, IL-33 signals through Myd88-dependent activation of the canonical NFκB and MAPK signalling pathways (20). The IL-33-induced signature in mast cells contained several proteins previously shown to be induced in LPS-stimulated macrophages, including Il6, Il1a, Il1b, Ptgs2, Nfkbiz, Junb, Tnfaip3, Nfkb2, and Traf1 (65-68). Notably IL-33 also induced components of the non-canonical NFκB system, as well as receptors able to activate this pathway. This may allow longer term induction of NFκB-dependent genes, however its relevance to mast cell biology is currently unclear and requires further investigation.

A subgroup of IL-33 induced proteins identified here are not typically associated with TLR-Myd88 signalling in macrophages, suggesting that they may represent a more mast cell-specific signature. These include Il9, Il13, Tph1, Hdc, Ntrk1, Tpbg, Nabp1, Hacd3, Tmem63b, Dennd5a, Herc1, and Cish. Of these, Il9 and Il13 are Th2-promoting cytokines, and their production is consistent with the established role of mast cells in Th2-driven immune responses (1-3). Similarly, Hdc and Tph1 are required for the synthesis of the biogenic amines histamine and serotonin, respectively (4).

Cish is related to the SOCS family and is known to be induced by cytokines that activate STAT5 signalling, as well as by IL-4-mediated STAT6 signalling. Once induced, Cish can inhibit signalling downstream of these cytokines (69). As mast cells are responsive to IL-3 and IL-4, it is therefore possible that IL-33 stimulation may reduce mast cell sensitivity to these cytokines. Notably, Cish knockout mice have been reported to develop a spontaneous allergic pulmonary phenotype and to display increased sensitivity in models of allergic asthma (70).

Mast cells are known to be involved in neuroimmune signalling (71). Relevant to this mast cells have been shown to both produce and respond to NGF (Neuronal Growth Factor) (72). We found that all three mast cell culture systems upregulated Ntrk1, the high affinity NGF receptor, suggesting that IL-33 may prime mast cells to respond to NGF, which can act as a chemoattractant for mast cells (73, 74).

FLMCs, BMMCs and PMCs also upregulated the cholesterol transport protein Gramd1a, while FLMCs and BMMCs also upregulated the related protein Gramd1b. In cells, membrane cholesterol is found at highest levels in the plasma membrane while cholesterol levels are sensed in the endoplasmic reticulum, which determines if the cell needs to activate cholesterol biosynthesis or uptake (75, 76). Gramd1 is involved in cholesterol sensing via transporting cholesterol from the plasma membrane to the endoplasmic reticulum (76), and increased levels of Gramd1a have been reported to result in higher levels of cholesterol biosynthetic enzymes in HTC116 cells (77). Related to this, we did find that the rate limiting enzyme in cholesterol biosynthesis, Hmgcr, was also increased by IL-33 in mast cells, although the fold increase did not pass the fold change cutoff used in FLMCs. Hmgcr activity has been reported to affect mast cell activity, as blocking is activity with statins has been shown to reduce degranulation downstream of the IgE receptor and IL-6 and TNF production downstream of IL-33 (76, 78, 79). A similar effect on Hmgcr expression downstream of Myd88-dependent signalling has also been reported in B cells (80). The effect in macrophages is more complex; TLRs that do not induce type I interferon activate Hmgcr expression while those which do, such as TLR3 and TLR4 repress Hmgcr expression via an autocrine interferon feedback loop (75).

It is likely that multiple pathways will regulate changes in protein abundance downstream of IL-33, and that these could act at multiple points including transcription, mRNA stability, translation and protein stability. Several studies have reported transcriptional analysis of IL-33 induced changes in mast cells (50, 81, 82). While these overlap with the changes detected here at the protein level, this was not complete (data not shown). It is however hard to extrapolate from transcriptomic to proteomic data, as frequently mRNA induction can be transient and will precede changes in protein level.

A strong signature for targets of the canonical NFκB pathway was observed in the upregulated proteins and this pathway is known to be activated by IL-33. Previous studies have also found that p38 inhibitors are effective at blocking IL-33 induced cytokine production in BMMCs, as well as type 2 innate lymphoid cells and dendritic cells (15, 16, 83, 84). We extended this to establish that p38 was also promoted the expression of a subset of IL-33 induced proteins in BMMCs, and that the effect of blocking p38 was greater than blocking the classical MAPKs ERK1 and ERK2. How p38 regulates these proteins likely involves multiple mechanisms. In macrophages, p38 controls both transcription and mRNA stability/translation downstream of Myd88-dependent signalling (15, 56, 58), and it is probable that both mechanisms also occur in mast cells. In macrophages extensive work has characterised a p38-MAPK activated protein kinase 2 (MK2)-Zfp36 axis that controls the mRNA stability and/or translation of mRNAs containing AU-rich elements in their 3’ UTR (reviewed in (58)). mRNAs containing AU rich elements were enriched in those that encode for IL-33 upregulated relative to down regulated proteins, and multiple p38 targets downstream of IL-33 in BMMCs contained AU rich elements in their mRNA. The proteomics dataset indicated that mast cells did not express Zfp36, which agrees with previous genetic evidence indicating that Zfp36 does not play a role in these cells (85). The expression of Zfp36l1 and Zfp36l2 was however detected in mast cells and these are likely to substitute for Zfp36 downstream of p38-MK2 signalling in IL-33 stimulated mast cells (15). Other proteins have also been implicated in regulation mRNA stability is mast cells including Zc3h12a and Zc3h12c (also known as Regnase) which has been shown to regulate mRNA stability downstream of IgE receptor stimulation in BMMCs (86). Regnase activity has further been shown to be controlled by IKK downstream of Myd88 in IL-1 stimulated fibroblasts (87) and IL-33 stimulated ILC2s (88). Thus downstream of IL-33, both p38 and IKK activity may affect the stability and/or translation of mRNAs containing AU rich elements in mast cells. Further work will be needed to resolve the different molecular mechanisms downstream of p38 controlling protein abundance.

Overall, our data provide a comprehensive proteomic resource for murine mast cell culture systems and demonstrate that, despite sharing core mast cell features, PMCs, BMMCs and FLMCs differ substantially in their maturation state and protease expression profiles. We further define a conserved IL-33 induced protein signature enriched for NFκB-, and p38-dependent pathways, Collectively, these findings extend previous transcriptomic studies by defining the proteomic landscape downstream of IL-33 in mast cells and provide a framework for future studies investigating how IL-33 regulates mast cell function in inflammation and allergic disease.

## Methods

### Animals

C57Bl6/J mice were obtained Charles River Laboratories or bred in house. Animals were kept in individually ventilated cages is specific pathogen free conditions. Mice were provided with free access to food (R&M3) and water. Animal rooms were on a 12/12 h light/dark cycle and maintained at 21 °C and between 45 and 65% humidity. Mice were sacrificed using an increasing concentration of CO2 and death confirmed by cervical dislocation. For the isolation of embryos, breeding pairs were set up for 1 week and plug checked daily. Embryos were isolated at E15.5. Work was carried out in accordance with UK law and subject to local ethical review by the University of Dundee Welfare and Ethical Use of Animals Committee. All work was carried at the University of Dundee out under a UK Home Office licence (PAAE38C7B) and approved by the University of Dundee Ethical Review and Welfare Committee.

### Foetal Liver Derived Mast Cell (FLMC) culture

Foetal livers were isolated from mouse embryos at E15.5 and disaggregated in Dulbecco’s phosphate buffered saline (D-PBS) by passing through a 70 µm cell strainer to obtain single cell suspensions. Suspensions were then centrifuged at 300*g* for 5 min at 21 °C and pellets were resuspended in 25 ml FLMC media (RPMI (Gibco) supplemented with 15% (v/v) FBS (Labtech), 4 mM L-glutamine (Gibco), 100 U/ml Penicillin, 100 µg/ml Streptomycin (Gibco), 25 mM HEPES pH7.5 (Gibco), 1 mM Sodium pyruvate (Gibco), 100 μM NEAA (Gibco), 50 μM β-mercaptoethanol (Sigma Aldrich), 10 ng/ml IL-3 (Peprotech), 30 ng/ml stem cell factor (SCF) (Peprotech)) and cultured at 37 °C, 5% CO_2_. Adherent cells were removed from culture and suspension cells were passaged bi-weekly with fresh FLMC media for up to 8 weeks, maintaining the cultures at approximately 1×10^6^ ml^-1^. Cells were used for experimentation 2 days after being passaged from week 6 onwards.

### Bone Marrow Derived Mast Cell (BMMC) culture

Bone marrow was flushed from the femurs and tibiae of adult mice using sterile D-PBS. The bone marrow suspension was passed through a 100 μm cell strainer and then centrifuged at 149*g* for 5 min. The supernatant was aspirated, and the cell pellet was resuspended in 25 ml of BMMC media (RPMI (Gibco) supplemented with 10% (v/v) FBS (Labtech), 5 mM L-glutamine (Gibco), 100 U/ml Penicillin, 100 µg/ml Streptomycin (Gibco), 25 mM HEPES pH7.5 (Gibco), 1 mM Sodium pyruvate (Gibco), 50 μM Non-Essential Amino Acids (NEAA) (Gibco), 50 μM β-mercaptoethanol (Sigma Aldrich), 30 ng/ml IL-3 (Peprotech)) and cultured at 37 °C, 5% CO_2_. Adherent cells were removed from culture and suspension cells were passaged bi-weekly with fresh BMMC media for up to 8 weeks, maintaining the cultures at approximately 1×10^6^ ml^-1^. Cells were used for experiments 2 days after being passaged from week 6 onwards.

### Peritoneal Mast Cell (PMC) Culture

The peritoneal cavity of adult mice was washed with approximately 5 ml D-PBS supplemented with 2 mM EDTA and 0.5% (w/v) BSA. Peritoneal washes were centrifuged at 149*g* for 5 min, and the cell pellet was resuspended in 5 ml of PMC media (RPMI (Gibco), 10% (v/v) FBS (Labtech), 5 mM L-glutamine (Gibco), 100 U/ml Penicillin 100 µg/ml Streptomycin (Gibco), 25 mM HEPES pH7.5 (Gibco), 1 mM Sodium pyruvate (Gibco), 50 μM NEAA (Gibco), 50 μM β-mercaptoethanol (Sigma Aldrich), 30 ng/ml IL-3 (Peprotech), 20 ng/ml SCF (Peprotech)). After 24 h in culture, non-adherent cells were removed from culture and discarded. The remaining adherent cells received 5 ml fresh PMC media and were returned to the incubator. Suspension and adherent cells were passaged bi-weekly with fresh PMC media for 2 weeks, maintaining the cultures at approximately 1×10^6^ ml^-1^. Cells were used for experimentation 2 days after being passaged on day 14.

### Cell stimulations

Cells were stimulated with 10 ng/ml IL-33 (Preprotech) for 24 h for proteomic experiments. When required, cells were pre-incubated in 2 µM PD184352 (Selleck) or 1 µM VX745 (Selleck) or an equivalent concentration of DMSO for 1 h before stimulation.

### Mass spectrometry

Preparation of samples for proteomic analysis was carried out as described(89, 90). Briefly mast cells were pelleted, washed twice in PBS and lysed in 5% (w/v) SDS (Sigma Aldrich), 10 mM Tris (2-carboxyethyl)phosphine (TCEP) (Thermo Fisher Scientific) and 50 mM Triethylamonium bicarbonate (TEAB) (Fisher Scientific). Lysates were then incubated at 95 ^0^C for 5 min, sonicated and 1 μl Benzonase was added to degrade DNA and lysates incubated for 15 min at 37 °C. Proteins were alkylated, acidified, loaded onto mini S-Trap column (Profiti) and digested using Trypsin Gold (Promega) as detailed in (89)

1.5 μg of peptide was injected onto a nanoscale C18 reverse-phase chromatography system (UltiMate 3000 RSLC nano, Thermo Fisher Scientific). Samples were loaded onto a trap column (100 μm × 2 cm, PepMap nanoViper C18 column, 5 μm, 100 Å, Thermo Fisher Scientific) at 10 μl/min and equilibrated in 0.1% trifluoroacetic acid (TFA). The column was then washed at 10 μl/min with 0.1% TFA for 3 min and replaced in-line with a resolving C18 column (75 μm × 50 cm, PepMap RSLC C18 column, 2 μm, 100 Å, Thermo Fisher Scientific). Peptides were eluted from the column at a constant flow rate of 300 nl/minute with a line (89)ar gradient from 97% buffer A (0.1% Formic acid) / 3% elution buffer B (80% Acetontrile, 0.1% Formic acid) to 6% elution buffer B in 5 min, then from 6% elution buffer B to 35% elution buffer B in 115 min, and finally to 80% elution buffer B within 7 min. Eluted peptides were analysed on an Orbitrap Exploris 480 Mass Spectrometer. Data acquisition was initiated with a full MS survey scan using an easy spray source in positive mode with a spray voltage of 2.650 kV, MS1 Orbitrap resolution of 120,000 MS2 Orbitrap resolution of 30,000 and cycle frequency of 350-1,650 m/z.

Following the MS survey scan, a MS/MS DIA scan events was completed with collision energy mode in stepped, collision energy type set to normalised, HCD collision energies of 25.5%, 27%, 30%, Orbitrap resolution of 30,000, first Mass of 200, RF lens of 40%, AGC target set to Custom, normalised AGC target of 3,000% and maximum injection time mode set to custom with a time of 55 ms. Mass accuracy was confirmed prior to initiation of sample analysis and all data, in both the MS survey scan and the MS/MS DIA, was acquired in profile mode.

Spectronaut 19 was used to search all mass spectrometry spectra acquired in this project, using the directDIA option against the Mouse C57BL6J UniProt database (UP000000589, 20210316). The search parameters used are detailed in (90).

Protein intensity data acquired in Spectronaut was exported to Perseus (91), where protein copy number and concentration were calculated using the histone ruler method (92). Details of the full data set are included in Supplementary table 4, and raw mass spectrometry data will be made available in PRIDE on publication.

For proteomic data sets, significance was determined by unpaired t-tests on log transformed data. Where indicated individual proteins were further analyses by ANOVA, with details of F and p values given in Supplementary table 5. Heat maps were generated using Morpheus software (Broad Institute). Enrichment analysis was carried out using a background list of the proteins present in the relevant proteomic dataset using the DAVID bioinformatics server (93).

## Acknowledgements

We would like to thank the Fingerprints proteomics facility at the University of Dundee for mass spectrometry and Dr Andy Howden (University of Dundee) for technical advice.

## Author contributions

Data acquisition: M.C.S, I.R.P, N.J.D.

Data Analysis: M.C.S, N.J.D, J.S.C.A.

Manuscript preparation: M.C.S, N.J.D, J.S.C.A.

## References

1. Metcalfe, D. D., Baram, D., and Mekori, Y. A. (1997) Mast cells Physiol Rev 77, 1033–1079

2. Galli, S. J. (2000) Mast cells and basophils Curr Opin Hematol 7, 32–39

3. Galli, S. J., Borregaard, N., and Wynn, T. A. (2011) Phenotypic and functional plasticity of cells of innate immunity: macrophages, mast cells and neutrophils Nat Immunol 12, 1035–1044

4. Wernersson, S., and Pejler, G. (2014) Mast cell secretory granules: armed for battle Nat Rev Immunol 14, 478–494

5. Galli, S. J., and Tsai, M. (2012) IgE and mast cells in allergic disease Nat Med 18, 693–704

6. Kinet, J. P. (1999) The high-affinity IgE receptor (Fc epsilon RI): from physiology to pathology Annu Rev Immunol 17, 931–972

7. Kolkhir, P., Elieh-Ali-Komi, D., Metz, M., Siebenhaar, F., and Maurer, M. (2022) Understanding human mast cells: lesson from therapies for allergic and non-allergic diseases Nat Rev Immunol 22, 294–308

8. Pahima, H. T., and Dwyer, D. F. (2025) Update on mast cell biology J Allergy Clin Immunol 155, 1115–1123

9. McCurdy, J. D., Lin, T. J., and Marshall, J. S. (2001) Toll-like receptor 4-mediated activation of murine mast cells J Leukoc Biol 70, 977–984

10. Allakhverdi, Z., Smith, D. E., Comeau, M. R., and Delespesse, G. (2007) Cutting edge: The ST2 ligand IL-33 potently activates and drives maturation of human mast cells J Immunol 179, 2051–2054

11. Ho, L. H., Ohno, T., Oboki, K., Kajiwara, N., Suto, H., Iikura, M. et al. (2007) IL-33 induces IL-13 production by mouse mast cells independently of IgE-FcepsilonRI signals J Leukoc Biol 82, 1481–1490

12. Moulin, D., Donze, O., Talabot-Ayer, D., Mezin, F., Palmer, G., and Gabay, C. (2007) Interleukin (IL)-33 induces the release of pro-inflammatory mediators by mast cells Cytokine 40, 216–225

13. Mukai, K., Tsai, M., Saito, H., and Galli, S. J. (2018) Mast cells as sources of cytokines, chemokines, and growth factors Immunol Rev 282, 121–150

14. Theoharides, T. C., Kempuraj, D., Tagen, M., Conti, P., and Kalogeromitros, D. (2007) Differential release of mast cell mediators and the pathogenesis of inflammation Immunol Rev 217, 65–78

15. McCarthy, P. C., Phair, I. R., Greger, C., Pardali, K., McGuire, V. A., Clark, A. R. et al. (2019) IL-33 regulates cytokine production and neutrophil recruitment via the p38 MAPK-activated kinases MK2/3 Immunol Cell Biol 97, 54–71

16. Drube, S., Kraft, F., Dudeck, J., Muller, A. L., Weber, F., Gopfert, C. et al. (2016) MK2/3 Are Pivotal for IL-33-Induced and Mast Cell-Dependent Leukocyte Recruitment and the Resulting Skin Inflammation J Immunol 197, 3662–3668

17. Darling, N. J., Arthur, J. S. C., and Cohen, P. (2021) Salt-inducible kinases are required for the IL-33-dependent secretion of cytokines and chemokines in mast cells J Biol Chem 296, 100428

18. Scott, I. C., Majithiya, J. B., Sanden, C., Thornton, P., Sanders, P. N., Moore, T. et al. (2018) Interleukin-33 is activated by allergen- and necrosis-associated proteolytic activities to regulate its alarmin activity during epithelial damage Sci Rep 8, 3363

19. Carriere, V., Roussel, L., Ortega, N., Lacorre, D. A., Americh, L., Aguilar, L. et al. (2007) IL-33, the IL-1-like cytokine ligand for ST2 receptor, is a chromatin-associated nuclear factor in vivo Proc Natl Acad Sci U S A 104, 282–287

20. Schmitz, J., Owyang, A., Oldham, E., Song, Y., Murphy, E., McClanahan, T. K. et al. (2005) IL-33, an interleukin-1-like cytokine that signals via the IL-1 receptor-related protein ST2 and induces T helper type 2-associated cytokines Immunity 23, 479–490

21. McSorley, H. J., and Smyth, D. J. (2021) IL-33: A central cytokine in helminth infections Semin Immunol 53, 101532

22. Cayrol, C., and Girard, J. P. (2018) Interleukin-33 (IL-33): A nuclear cytokine from the IL-1 family Immunol Rev 281, 154–168

23. Griesenauer, B., and Paczesny, S. (2017) The ST2/IL-33 Axis in Immune Cells during Inflammatory Diseases Front Immunol 8, 475

24. Moritz, D. R., Rodewald, H. R., Gheyselinck, J., and Klemenz, R. (1998) The IL-1 receptor-related T1 antigen is expressed on immature and mature mast cells and on fetal blood mast cell progenitors J Immunol 161, 4866–4874

25. Joulia, R., L’Faqihi, F. E., Valitutti, S., and Espinosa, E. (2017) IL-33 fine tunes mast cell degranulation and chemokine production at the single-cell level J Allergy Clin Immunol 140, 497–509 e410

26. Franke, K., Wang, Z., Zuberbier, T., and Babina, M. (2021) Cytokines Stimulated by IL-33 in Human Skin Mast Cells: Involvement of NF-kappaB and p38 at Distinct Levels and Potent Co-Operation with FcepsilonRI and MRGPRX2 Int J Mol Sci 22,

27. Wang, Z., Guhl, S., Franke, K., Artuc, M., Zuberbier, T., and Babina, M. (2019) IL-33 and MRGPRX2-Triggered Activation of Human Skin Mast Cells-Elimination of Receptor Expression on Chronic Exposure, but Reinforced Degranulation on Acute Priming Cells 8,

28. Babina, M., Wang, Z., Franke, K., Guhl, S., Artuc, M., and Zuberbier, T. (2019) Yin-Yang of IL-33 in Human Skin Mast Cells: Reduced Degranulation, but Augmented Histamine Synthesis through p38 Activation J Invest Dermatol 139, 1516–1525 e1513

29. Ronnberg, E., Ghaib, A., Ceriol, C., Enoksson, M., Arock, M., Safholm, J. et al. (2019) Divergent Effects of Acute and Prolonged Interleukin 33 Exposure on Mast Cell IgE-Mediated Functions Front Immunol 10, 1361

30. Wang, J. X., Kaieda, S., Ameri, S., Fishgal, N., Dwyer, D., Dellinger, A. et al. (2014) IL-33/ST2 axis promotes mast cell survival via BCLXL Proc Natl Acad Sci U S A 111, 10281–10286

31. Bradding, P., and Pejler, G. (2018) The controversial role of mast cells in fibrosis Immunol Rev 282, 198–231

32. Varricchi, G., Marone, G., and Kovanen, P. T. (2020) Cardiac Mast Cells: Underappreciated Immune Cells in Cardiovascular Homeostasis and Disease Trends Immunol 41, 734–746

33. Tauber, M., Basso, L., Martin, J., Bostan, L., Pinto, M. M., Thierry, G. R. et al. (2023) Landscape of mast cell populations across organs in mice and humans J Exp Med 220,

34. Iuliano, C., Absmaier-Kijak, M., Sinnberg, T., Hoffard, N., Hils, M., Koberle, M. et al. (2022) Fetal Tissue-Derived Mast Cells (MC) as Experimental Surrogate for In Vivo Connective Tissue MC Cells 11,

35. Tsvilovskyy, V., Solis-Lopez, A., Ohlenschlager, K., and Freichel, M. (2018) Isolation of Peritoneum-derived Mast Cells and Their Functional Characterization with Ca2+-imaging and Degranulation Assays J Vis Exp 10.3791/57222

36. McNeil, B. D., Pundir, P., Meeker, S., Han, L., Undem, B. J., Kulka, M. et al. (2015) Identification of a mast-cell-specific receptor crucial for pseudo-allergic drug reactions Nature 519, 237–241

37. Green, D. P., Limjunyawong, N., Gour, N., Pundir, P., and Dong, X. (2019) A Mast-Cell-Specific Receptor Mediates Neurogenic Inflammation and Pain Neuron 101, 412–420 e413

38. Dwyer, D. F., Barrett, N. A., Austen, K. F., and Immunological Genome Project, C. (2016) Expression profiling of constitutive mast cells reveals a unique identity within the immune system Nat Immunol 17, 878–887

39. Fukuishi, N., Murakami, S., Ohno, A., Yamanaka, N., Matsui, N., Fukutsuji, K. et al. (2014) Does beta-hexosaminidase function only as a degranulation indicator in mast cells? The primary role of beta-hexosaminidase in mast cell granules J Immunol 193, 1886–1894

40. Orange, R. P., and Moore, E. G. (1976) Functional characterization of rat mast cell arylsulfatase activity J Immunol 117, 2191–2196

41. Aubert, A., Jung, K., Hiroyasu, S., Pardo, J., and Granville, D. J. (2024) Granzyme serine proteases in inflammation and rheumatic diseases Nat Rev Rheumatol 20, 361–376

42. Phair, I. R., Sumoreeah, M. C., Scott, N., Spinelli, L., and Arthur, J. S. C. (2022) IL-33 induces granzyme C expression in murine mast cells via an MSK1/2-CREB-dependent pathway Biosci Rep 42,

43. Strik, M. C., de Koning, P. J., Kleijmeer, M. J., Bladergroen, B. A., Wolbink, A. M., Griffith, J. M. et al. (2007) Human mast cells produce and release the cytotoxic lymphocyte associated protease granzyme B upon activation Mol Immunol 44, 3462–3472

44. Abrink, M., Grujic, M., and Pejler, G. (2004) Serglycin is essential for maturation of mast cell secretory granule J Biol Chem 279, 40897–40905

45. Henningsson, F., Hergeth, S., Cortelius, R., Abrink, M., and Pejler, G. (2006) A role for serglycin proteoglycan in granular retention and processing of mast cell secretory granule components FEBS J 273, 4901–4912

46. Chen, C. Y., Lee, J. B., Liu, B., Ohta, S., Wang, P. Y., Kartashov, A. V. et al. (2015) Induction of Interleukin-9-Producing Mucosal Mast Cells Promotes Susceptibility to IgE-Mediated Experimental Food Allergy Immunity 43, 788–802

47. Medzhitov, R., Preston-Hurlburt, P., Kopp, E., Stadlen, A., Chen, C., Ghosh, S. et al. (1998) MyD88 is an adaptor protein in the hToll/IL-1 receptor family signaling pathways Mol Cell 2, 253–258

48. Sun, S. C. (2017) The non-canonical NF-kappaB pathway in immunity and inflammation Nat Rev Immunol 17, 545–558

49. Vallabhapurapu, S., and Karin, M. (2009) Regulation and function of NF-kappaB transcription factors in the immune system Annu Rev Immunol 27, 693–733

50. Jordan, P. M., Andreas, N., Groth, M., Wegner, P., Weber, F., Jager, U. et al. (2021) ATP/IL-33-triggered hyperactivation of mast cells results in an amplified production of pro-inflammatory cytokines and eicosanoids Immunology 164, 541–554

51. Wajant, H. (2015) Principles of antibody-mediated TNF receptor activation Cell Death Differ 22, 1727–1741

52. Bain, J., Plater, L., Elliott, M., Shpiro, N., Hastie, C. J., McLauchlan, H. et al. (2007) The selectivity of protein kinase inhibitors: a further update Biochem J 408, 297–315

53. McGuire, V. A., Gray, A., Monk, C. E., Santos, S. G., Lee, K., Aubareda, A. et al. (2013) Cross talk between the Akt and p38alpha pathways in macrophages downstream of Toll-like receptor signaling Mol Cell Biol 33, 4152–4165

54. Huang, H., Li, Y., Liang, J., and Finkelman, F. D. (2018) Molecular Regulation of Histamine Synthesis Front Immunol 9, 1392

55. Yabut, J. M., Desjardins, E. M., Chan, E. J., Day, E. A., Leroux, J. M., Wang, B. et al. (2020) Genetic deletion of mast cell serotonin synthesis prevents the development of obesity and insulin resistance Nat Commun 11, 463

56. Arthur, J. S., and Ley, S. C. (2013) Mitogen-activated protein kinases in innate immunity Nat Rev Immunol 13, 679–692

57. Tiedje, C., Holtmann, H., and Gaestel, M. (2014) The role of mammalian MAPK signaling in regulation of cytokine mRNA stability and translation J Interferon Cytokine Res 34, 220–232

58. O’Neil, J. D., Ammit, A. J., and Clark, A. R. (2018) MAPK p38 regulates inflammatory gene expression via tristetraprolin: Doing good by stealth Int J Biochem Cell Biol 94, 6–9

59. Bakheet, T., Hitti, E., and Khabar, K. S. A. (2018) ARED-Plus: an updated and expanded database of AU-rich element-containing mRNAs and pre-mRNAs Nucleic Acids Res 46, D218–D220

60. Plum, T., Wang, X., Rettel, M., Krijgsveld, J., Feyerabend, T. B., and Rodewald, H. R. (2020) Human Mast Cell Proteome Reveals Unique Lineage, Putative Functions, and Structural Basis for Cell Ablation Immunity 52, 404–416 e405

61. Kovacs, D., Heger, K., Giansanti, P., Iuliano, C., Meissner, F., Mann, M. et al. (2025) Mast cells modulate macrophage biology through release of prestored CSF1 J Allergy Clin Immunol 156, 1260–1276

62. Gentek, R., Ghigo, C., Hoeffel, G., Bulle, M. J., Msallam, R., Gautier, G. et al. (2018) Hemogenic Endothelial Fate Mapping Reveals Dual Developmental Origin of Mast Cells Immunity 48, 1160–1171 e1165

63. Dai, H., and Korthuis, R. J. (2011) Mast Cell Proteases and Inflammation Drug Discov Today Dis Models 8, 47–55

64. Malbec, O., Roget, K., Schiffer, C., Iannascoli, B., Dumas, A. R., Arock, M. et al. (2007) Peritoneal cell-derived mast cells: an in vitro model of mature serosal-type mouse mast cells J Immunol 178, 6465–6475

65. He, L., Jhong, J. H., Chen, Q., Huang, K. Y., Strittmatter, K., Kreuzer, J. et al. (2021) Global characterization of macrophage polarization mechanisms and identification of M2-type polarization inhibitors Cell Rep 37, 109955

66. Yamamoto, M., Yamazaki, S., Uematsu, S., Sato, S., Hemmi, H., Hoshino, K. et al. (2004) Regulation of Toll/IL-1-receptor-mediated gene expression by the inducible nuclear protein IkappaBzeta Nature 430, 218–222

67. Bhatt, D. M., Pandya-Jones, A., Tong, A. J., Barozzi, I., Lissner, M. M., Natoli, G. et al. (2012) Transcript dynamics of proinflammatory genes revealed by sequence analysis of subcellular RNA fractions Cell 150, 279–290

68. Baker, C. P., Laba, S., Warner, J., Shepherd, K., Wilson, H. M., and Arthur, J. S. C. (2026) Candida albicans infection suppresses lipopolysaccharide or Pseudomonas aeruginosa stimulated murine bone marrow derived macrophage (BMDM) responses Sci Rep 16,

69. Naser, W., and Ward, A. C. (2026) Cytokine inducible SH2-containing protein: a versatile negative regulator of cytokine receptor signaling Front Immunol 17, 1752876

70. Yang, X. O., Zhang, H., Kim, B. S., Niu, X., Peng, J., Chen, Y. et al. (2013) The signaling suppressor CIS controls proallergic T cell development and allergic airway inflammation Nat Immunol 14, 732–740

71. Plum, T., Feyerabend, T. B., and Rodewald, H. R. (2024) Beyond classical immunity: Mast cells as signal converters between tissues and neurons Immunity 57, 2723–2736

72. Nilsson, G., Forsberg-Nilsson, K., Xiang, Z., Hallbook, F., Nilsson, K., and Metcalfe, D. D. (1997) Human mast cells express functional TrkA and are a source of nerve growth factor Eur J Immunol 27, 2295–2301

73. Sawada, J., Itakura, A., Tanaka, A., Furusaka, T., and Matsuda, H. (2000) Nerve growth factor functions as a chemoattractant for mast cells through both mitogen-activated protein kinase and phosphatidylinositol 3-kinase signaling pathways Blood 95, 2052–2058

74. Skaper, S. D. (2017) Nerve growth factor: a neuroimmune crosstalk mediator for all seasons Immunology 151, 1–15

75. Lee, M. S., and Bensinger, S. J. (2022) Reprogramming cholesterol metabolism in macrophages and its role in host defense against cholesterol-dependent cytolysins Cell Mol Immunol 19, 327–336

76. Naito, T., and Saheki, Y. (2021) GRAMD1-mediated accessible cholesterol sensing and transport Biochim Biophys Acta Mol Cell Biol Lipids 1866, 158957

77. Zhang, C., Yu, R., Li, S., Yuan, M., Hu, T., Liu, J. et al. (2025) KRAS mutation increases histone H3 lysine 9 lactylation (H3K9la) to promote colorectal cancer progression by facilitating cholesterol transporter GRAMD1A expression Cell Death Differ 32, 2225–2238

78. Sahid, M. N. A., Liu, S., Kiyoi, T., and Maeyama, K. (2017) Inhibition of the mevalonate pathway by simvastatin interferes with mast cell degranulation by disrupting the interaction between Rab27a and double C2 alpha proteins Eur J Pharmacol 814, 255–263

79. Kolawole, E. M., McLeod, J. J., Ndaw, V., Abebayehu, D., Barnstein, B. O., Faber, T. et al. (2016) Fluvastatin Suppresses Mast Cell and Basophil IgE Responses: Genotype-Dependent Effects J Immunol 196, 1461–1470

80. Cheung, D. M. S., Razsolkov, M., Bonacina, F., Andrews, S., Sumoreeah, M. C., Sinclair, L. V. et al. (2026) Lipopolysaccharide stimulates dynamic changes in B cell metabolism to promote proliferation Elife 14,

81. Li, Q., Chen, W., Sun, M., Ni, X., Ran, Q., Hu, X. et al. (2026) Stromal Cell-Mast Cell Communication Orchestrates Anti-Viral Immunity in the Meninges Adv Sci (Weinh) 13, e14842

82. Chhiba, K. D., Hsu, C. L., Berdnikovs, S., and Bryce, P. J. (2017) Transcriptional Heterogeneity of Mast Cells and Basophils upon Activation J Immunol 198, 4868–4878

83. Petrova, T., Pesic, J., Pardali, K., Gaestel, M., and Arthur, J. S. C. (2020) p38 MAPK signalling regulates cytokine production in IL-33 stimulated Type 2 Innate Lymphoid cells Sci Rep 10, 3479

84. Gopfert, C., Andreas, N., Weber, F., Hafner, N., Yakovleva, T., Gaestel, M. et al. (2018) The p38-MK2/3 Module Is Critical for IL-33-Induced Signaling and Cytokine Production in Dendritic Cells J Immunol 200, 1198–1206

85. Hochdorfer, T., Tiedje, C., Stumpo, D. J., Blackshear, P. J., Gaestel, M., and Huber, M. (2013) LPS-induced production of TNF-alpha and IL-6 in mast cells is dependent on p38 but independent of TTP Cell Signal 25, 1339–1347

86. Bataclan, M., Leoni, C., Moro, S. G., Pecoraro, M., Wong, E. H., Heissmeyer, V. et al. (2024) Crosstalk between Regnase-1 and -3 shapes mast cell survival and cytokine expression Life Sci Alliance 7,

87. Iwasaki, H., Takeuchi, O., Teraguchi, S., Matsushita, K., Uehata, T., Kuniyoshi, K. et al. (2011) The IkappaB kinase complex regulates the stability of cytokine-encoding mRNA induced by TLR-IL-1R by controlling degradation of regnase-1 Nat Immunol 12, 1167–1175

88. Matsushita, K., Tanaka, H., Yasuda, K., Adachi, T., Fukuoka, A., Akasaki, S. et al. (2020) Regnase-1 degradation is crucial for IL-33- and IL-25-mediated ILC2 activation JCI Insight 5,

89. Baker, C. P., Phair, I. R., Brenes, A. J., Atrih, A., Ryan, D. G., Bruderer, R. et al. (2022) DIA label-free proteomic analysis of murine bone-marrow-derived macrophages STAR Protoc 3, 101725

90. Baker, C. P., Bruderer, R., Abbott, J., Arthur, J. S. C., and Brenes, A. J. (2024) Optimizing Spectronaut Search Parameters to Improve Data Quality with Minimal Proteome Coverage Reductions in DIA Analyses of Heterogeneous Samples J Proteome Res 23, 1926–1936

91. Tyanova, S., Temu, T., Sinitcyn, P., Carlson, A., Hein, M. Y., Geiger, T. et al. (2016) The Perseus computational platform for comprehensive analysis of (prote)omics data Nat Methods 13, 731–740

92. Wisniewski, J. R., Hein, M. Y., Cox, J., and Mann, M. (2014) A “proteomic ruler” for protein copy number and concentration estimation without spike-in standards Mol Cell Proteomics 13, 3497–3506

93. Sherman, B. T., Panzade, G., Dotrang, T., Hao, M., Xu, L., Li, X. et al. (2026) DAVID: a web server for functional annotation and functional enrichment analysis of gene lists (2025 update) Nucleic Acids Res 10.1093/nar/gkag470

