## Supplementary figures 1-6 for "MAPK dependent IL-33 responses define a conserved inflammatory programme in mast cells"

Supplementary Figures 1- 7

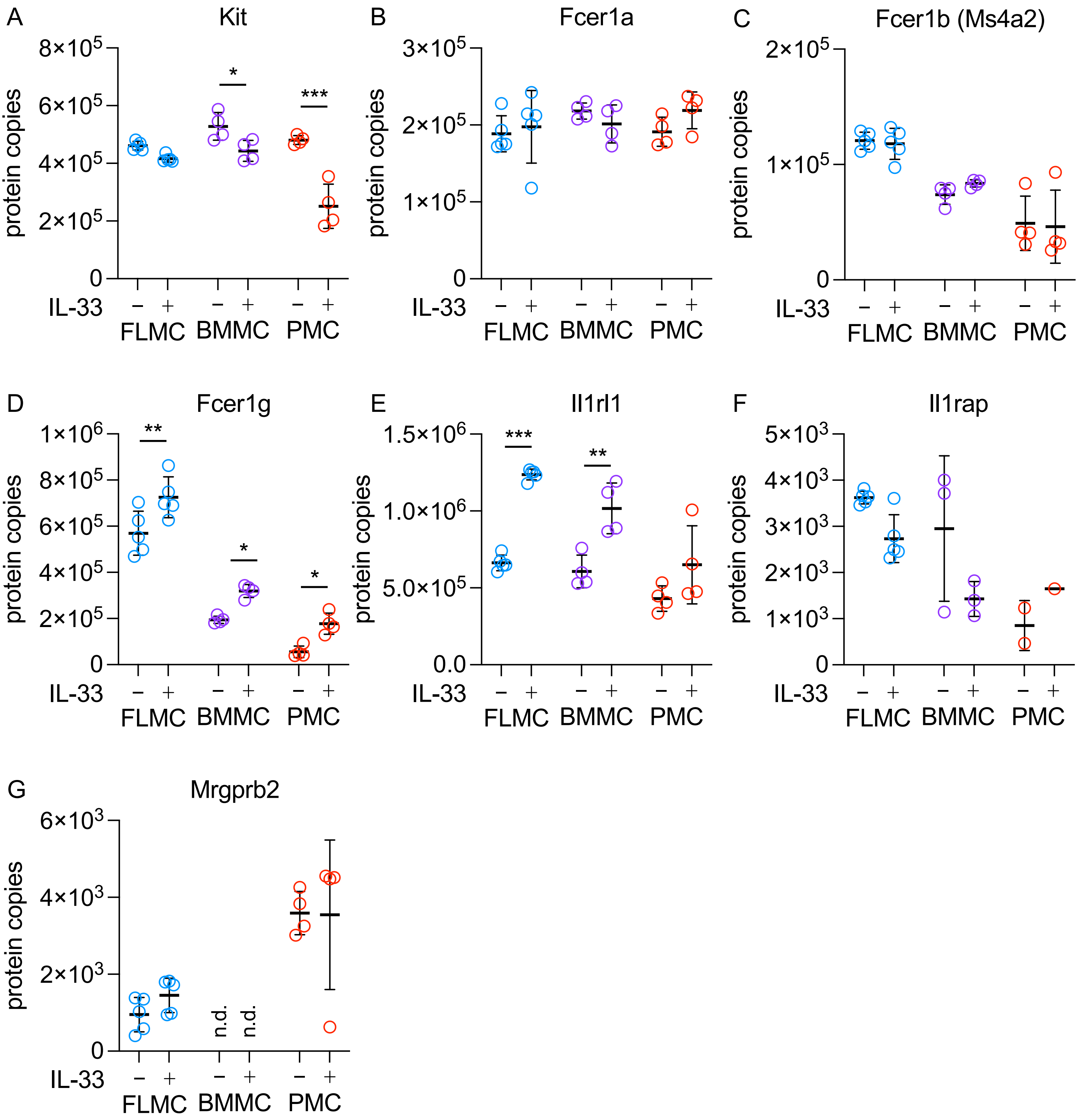


**Supplementary Figure 1. Expression of markers on cultured FLMCs, BMMCs and PMCs.**

FLMCs (n=5), BMMCs (n=4) and PMCs (n=4) were cultured as described in the methods. Their proteomes were determined with or without 10 ng/ml IL-33 stimulation for 24 h using DIA based mass spectrometry. Protein copy number was determined using the histone ruler method. The levels of Kit (A), Fcer1a (B), Fcer1b/Ms4a2 (C), Fcer1g (D), Il1rl1/ST2 (E), Il1rap (F) and Mrgprb2 (G) are shown. Graphs show mean and standard deviation with individual replicates shown as symbols; in F Il1rap was not detected in all replicates. Significance was determined by two way ANOVA with Sidak's multiple comparison tests to compare control and IL-33 in each cell type. *p*<0.05 is indicated by *, *p*<0.01 by ** and *p*<0.001 by ***. N.d. indicates not detected.

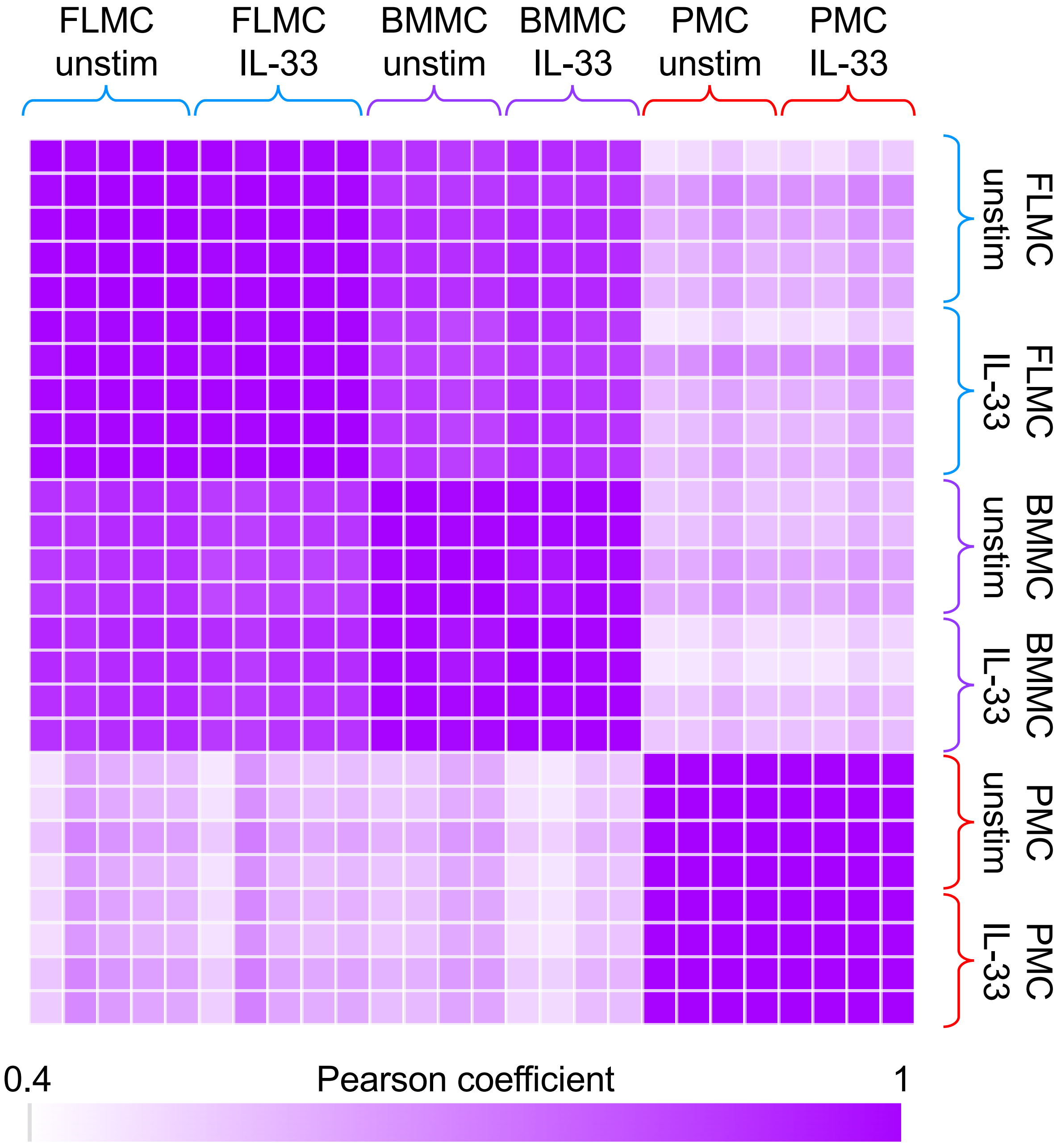


**Supplementary Figure 2. Mast cell classifications**.

FLMCs (n=5), BMMCs (n=4) and PMCs (n=4) were cultured as described in the methods. Their proteomes were determined with or without 10 ng/ml IL-33 stimulation for 24 h using DIA based mass spectrometry. A group of 159 proteins shown previously to be differentially expressed in connective tissue and mucosal mast cells was extracted from the proteomic data set. Pearson coefficients were determined for pairwise comparisons across all 26 replicates are shown in the heatmap.

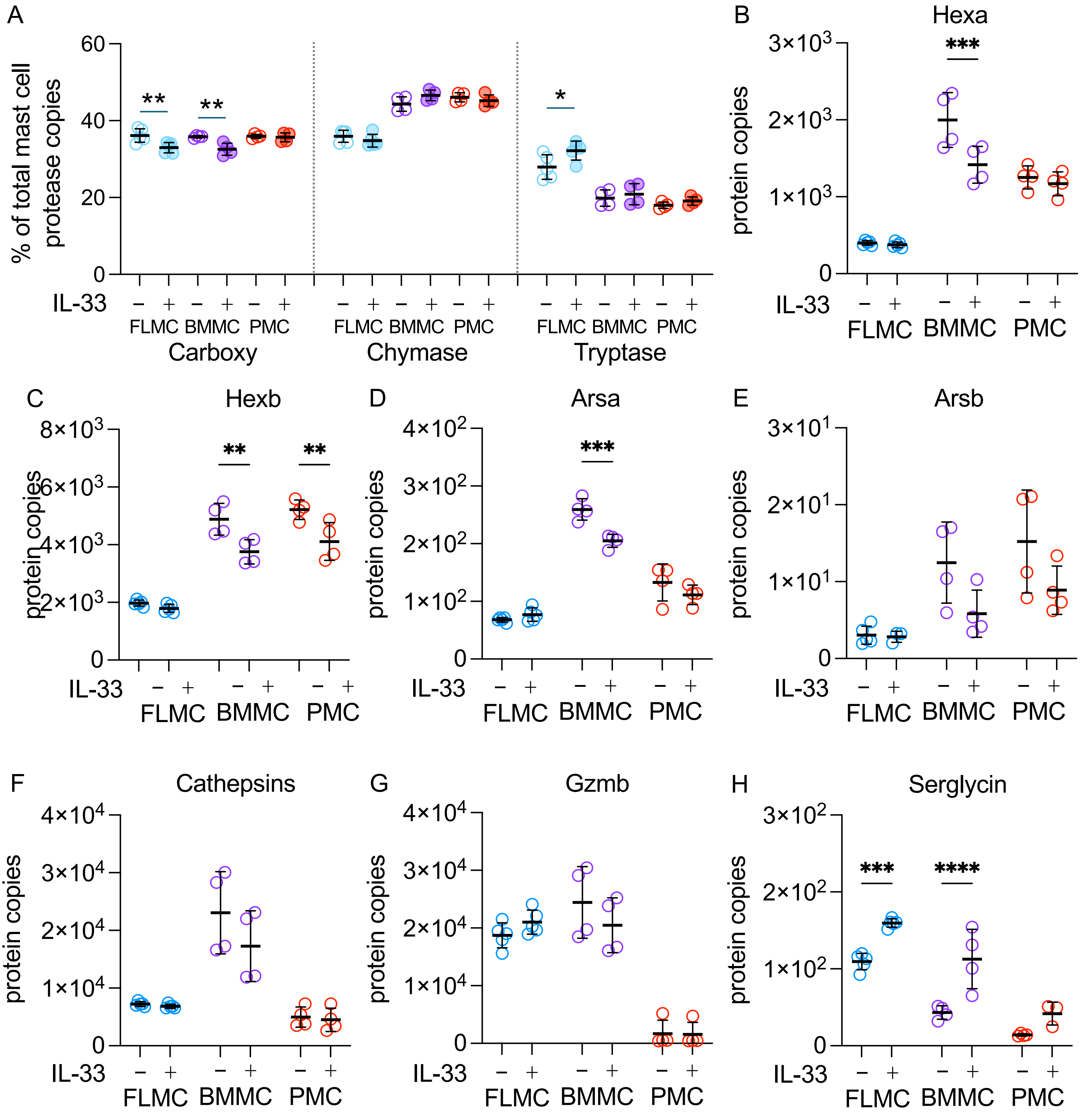


**Supplementary Figure 3. Abundance of granule proteins**.

FLMCs (n=5), BMMCs (n=4) and PMCs (n=4) were cultured as described in the methods. Their proteomes were determined with or without stimulation with 10 ng/ml IL-33 for 24 h using DIA based mass spectrometry. Protein copy number was estimated using the histone ruler method. The percentage of carboxypeptidase, chymases and tryptase is shown in (A). The levels of Hexa (B), Hexb (C), Arsa (D), Arsb (E), the sum of cathepsins B, C, D, E, G and L (F), Gzmb (G) and Srgn (H) are shown. Graphs show mean and standard deviation with individual replicates shown as symbols. Significance was determined by two way ANOVA with Sidak's multiple comparison tests to compare control and IL-33 in each cell type. $p<0.05$ is indicated by *, $p<0.01$ by ** and $p<0.001$ by ***.

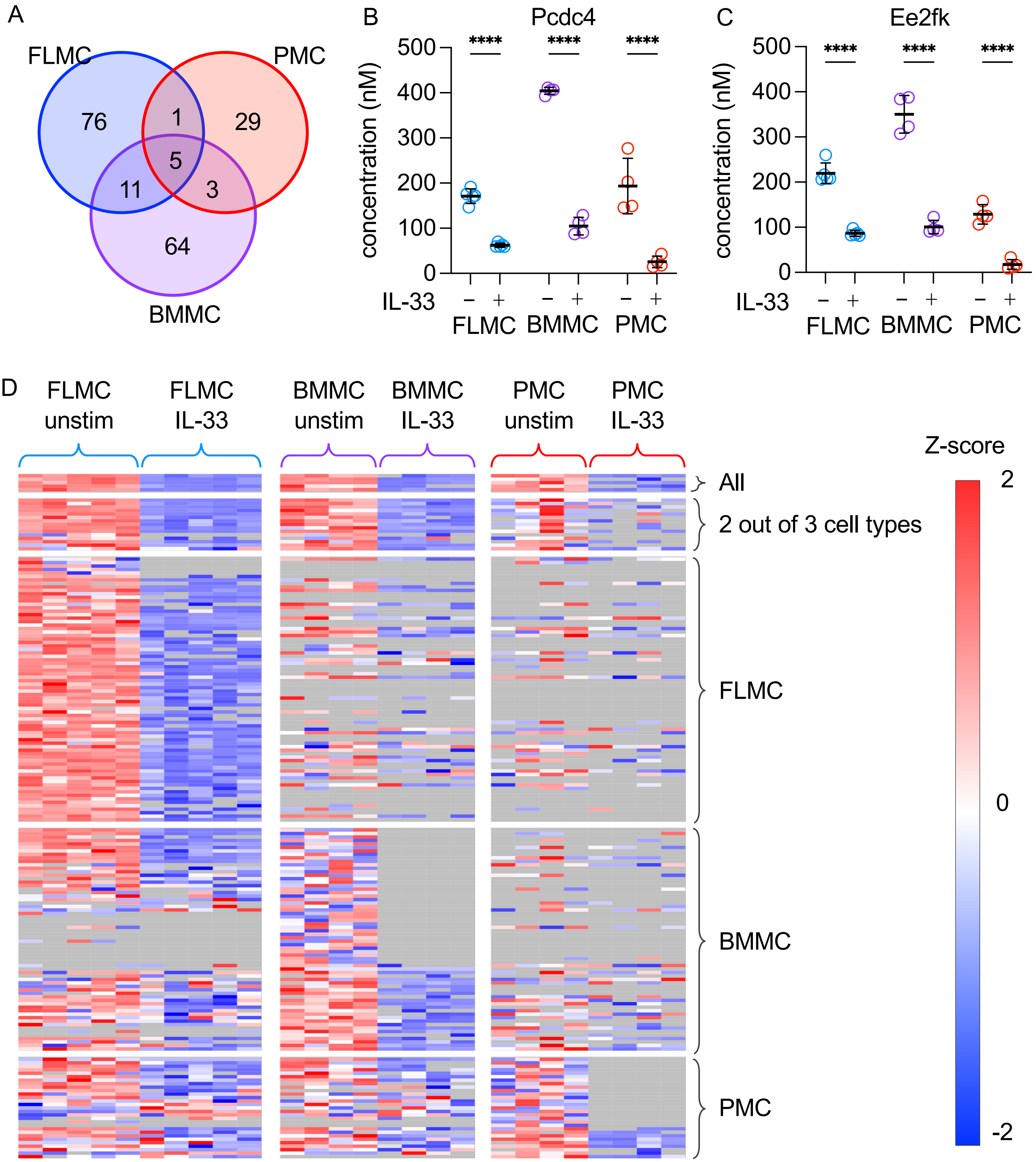


**Supplementary Figure 4. Proteins downregulated in response to IL-33 across different mast cell culture types.**

(A) Intersects between the proteins identified as downregulated in FLMCs, BMMCs and PMCs (based on a log2 fold change greater than 2 standard deviations from the median and $p < 0.01$ or present in no IL-33 replicates but all unstimulated replicates for a given cell type). 5 proteins (Espn, Sla, Pdcd4, Rcbtb2 and Eef2k) met these thresholds in all 3 cell types.
(B-C) Graphs show the estimated concentrations of Pcdc4 (B) and Ee2fk (C) in FLMCs, BMMCs and PMCs. For comparisons between unstimulated and IL-33 stimulated conditions, $p$<0.05 is indicated by *, $p$<0.01 by **, $p$<0.001 by *** and $p$<0.0001 by **** (Two-way ANOVA and Sidak's multiple comparison tests).
(D) Heat map of IL-33-regulated proteins in FLMCs, BMMCs and PMCs. Grey indicates replicate where a protein was not identified.

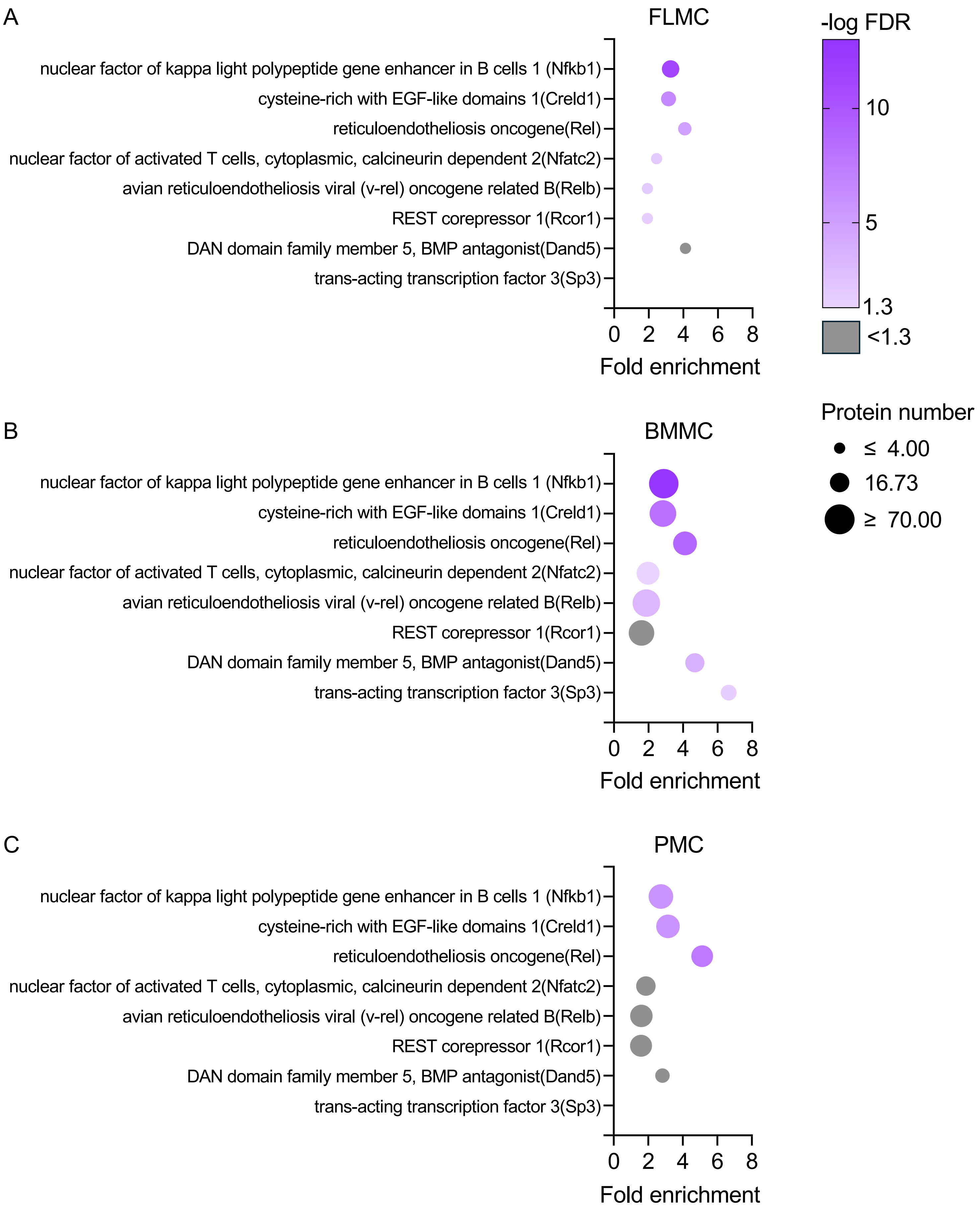


**Supplementary Figure 5. Transcription factor enrichment for IL-33 upregulated proteins**.

Upregulated protein list for FLMCs, BMMCs and PMCs were defined as in Figure 5. Enrichment was carried out against the TFLINK database in DAVID as described in the methods. Results for a binding site that passed a FDR cutoff of 0.05 for any of the 3 mast cell types are shown. Sp3 was not returned for enrichment analysis of FLMCs or PMCs.

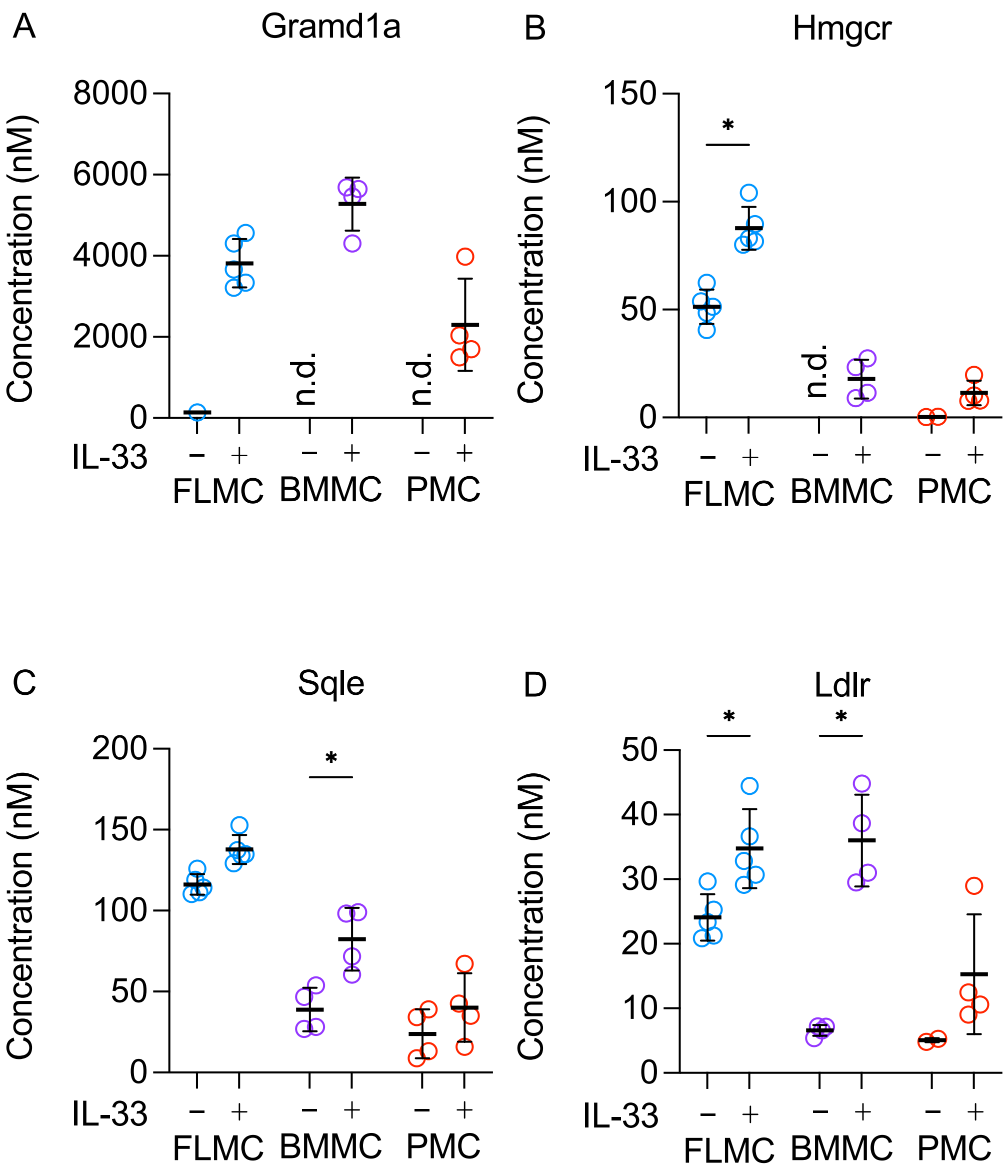


**Supplementary Figure 6. Abundance of enzymes involved in cholesterol metabolism.**

FLMCs (n=5), BMMCs (n=4) and PMCs (n=4) were cultured as described in the methods. Their proteomes were determined with or without stimulation with 10 ng/ml IL-33 for 24 h using DIA based mass spectrometry. Protein copy number was determined using the histone ruler method. The levels of Grand1a (A), Hmgcr (B), Sqle (C) and Ldlr (D) are shown. Graphs show mean and standard deviation with individual replicates shown as symbols. Significance was determined by two way ANOVA with Sidak's multiple comparison tests to compare control and IL-33 in each cell type. *p*<0.05 is indicated by *, *p*<0.01 by ** and *p*<0.001 by ***. N.d. indicates not detected.

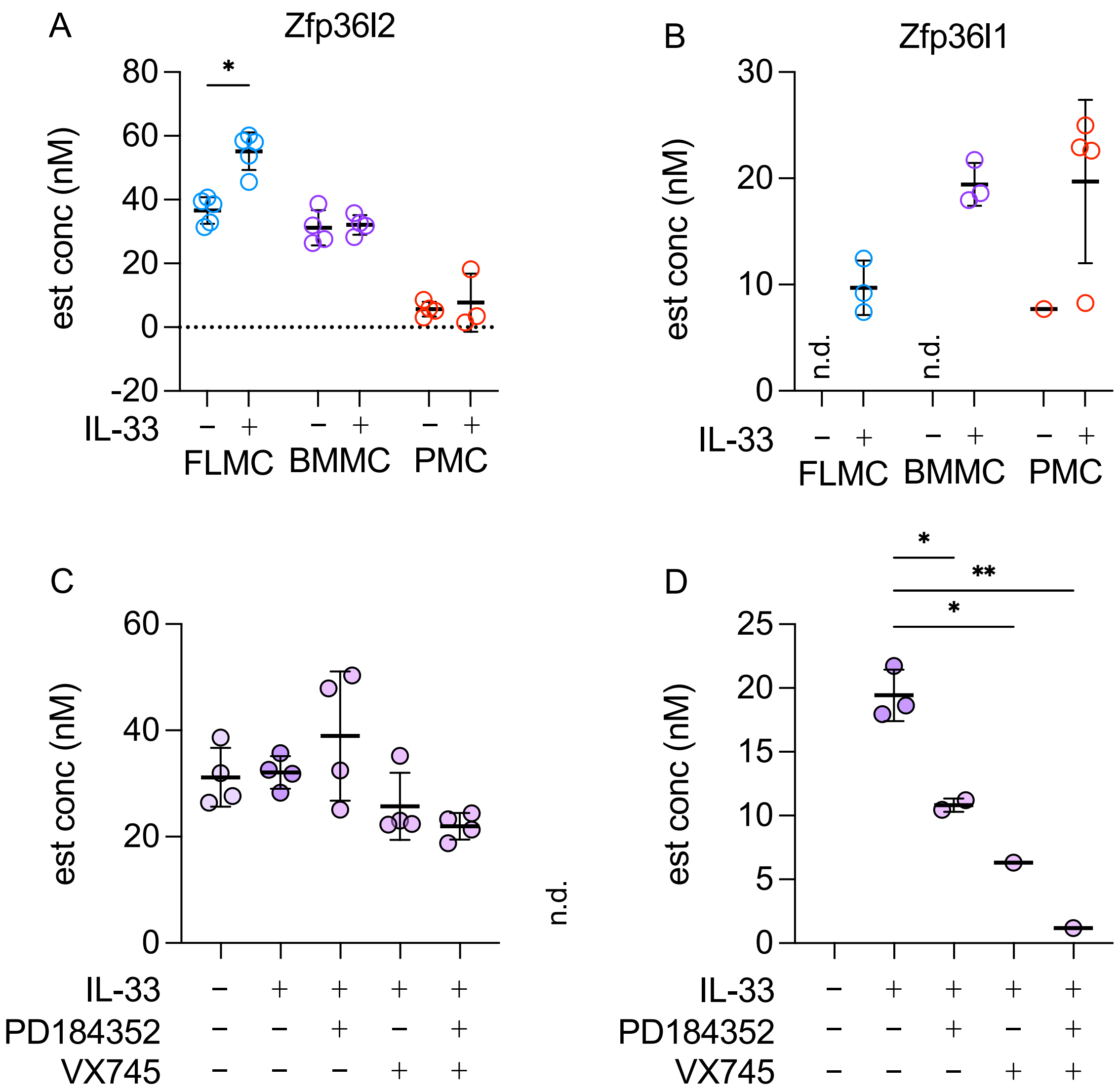


**Supplementary Figure 7. Abundance of Zfp36l2 and Zfp36l1 in mast cell proteomic datasets**.

The abundance of Zfp36l2 and Zfp36l1 in FLMCs, BMMCs and PMCs is shown in (A) and (B) respectively. * indicates $p$ <0.05 for comparisons between unstimulated and IL-33 treated cells (Two way ANOVA, F=8.196, $p$=0.0027 for the interaction, F=11.72, $p$=0.0029 for stimulation and F=116.97, $p$<0.0001 for cell type for Zfp36l2, test not valid for Zfp36l1 due to lack of detection in unstimulated samples).
The effect of either 2 µM PD184352, 1 µM VX745 or a combination of both inhibitors on IL-33 regulated protein levels in BMMCs is shown in (C) for Zfp36l2 and (D) for Zfp36l1. For comparisons to the IL-33 condition, $p$<0.05 is indicated by * and <0.01 by ** (One way ANOVA F=3.61, $p$=0.0297 for Zfp36l2 and F=37.63, $p$=0.0070 for Zfp36l1). Graphs show mean and standard deviation with symbol representing individual replicates. n.d. indicates the protein was not detected in any replicates for that condition.
